# Controlled linking of AAV capsids enables coordinated multi-vector delivery

**DOI:** 10.64898/2026.04.13.716639

**Authors:** Yongha Kim, Xiaozhe Ding, Chaeyun Yang, Gwangbin Lee, Viviana Gradinaru, Jimin Park

## Abstract

Adeno-associated virus (AAV)-based delivery of large genetic systems splits cargo across vectors, but functional reconstitution depends on stochastic co-delivery and often requires high viral doses. Here, we present a multi-capsid linking strategy that enables coordinated delivery of AAV vectors carrying distinct or split genetic cargos. Using a substrate-confined linking process, we suppress uncontrolled higher-order aggregation while enabling programmable nanoscale capsid coupling via oligonucleotide linkers. These linkers further enable efficient downstream purification of linked AAVs and recycling of unlinked vectors, yielding ∼70% AAV dimers and trimers in <24 hours. By enforcing coordinated delivery, linked AAVs achieve co-transduction at reduced viral doses and with greater expression uniformity than conventional dual-AAV approaches. This improved coordination enhances reconstitution of split genetic systems and increases targeted prime editing efficiency in vitro and in the mouse brain. This capsid coupling strategy expands the functional scope of AAVs toward dose-efficient and uniform delivery of large genetic systems.

## Introduction

Adeno-associated virus (AAV) vectors have been widely applied as in vivo gene delivery vehicles because of their low immunogenicity, tunable tropism, and ability to support long-term transgene expression^1–6^. However, their utility is fundamentally constrained by an intrinsic genome packaging capacity of ∼4.7 kilobases (kb). This size limit precludes delivery of many functional genetic cargos, including CRISPR-based genome editing systems and a substantial fraction of human cDNAs associated with genetic disease, restricting both experimental design and therapeutic scope^7–13^.

The most common strategy to address this limitation is the split- or dual-AAV approach, in which an oversized genetic cargo is divided across multiple vectors and reconstituted within the target cell^14–17^. Using this strategy, functional delivery has been demonstrated for diverse classes of cargos, including genome editors and large therapeutic protein-coding genes^7–13^. However, functional reconstitution relies on stochastic co-delivery of independent vectors, resulting in intrinsically low co-transduction efficiency and pronounced cell-to-cell heterogeneity. In practice, achieving robust functional outcomes often requires high viral doses, raising concerns related to toxicity, immune responses, and variability in in vivo settings^18–20^.

To address this challenge, pre-linking multiple AAV capsids prior to delivery has been proposed as a means to enforce coordinated co-entry of split genetic cargos. For example, AAV multimers and AAV-functional material conjugates have been generated using protein-based capsid linkers or chemical conjugation motifs^21, 22^. However, these strategies are typically implemented in homogeneous solutions, where uncontrolled chain reactions generate broad distributions of linked products. Although downstream purification with high-performance liquid chromatography or ultracentrifugation can enrich for defined species such as dimers or trimers, this is generally achieved at the expense of yield and process simplicity^21^. As a result, scalable production strategies for well-defined linked AAVs have not been established.

Notably, the field of nanomaterials engineering has developed a broad set of toolkits that enable precise control over multimer formation^23–26^. A unifying feature of these approaches is stepwise assembly of nanoparticles onto solid substrates, which confines assembly to interfaces and limits uncontrolled solution-phase reactions. In these approaches, nucleic-acid linkers are commonly introduced onto nanoparticle surfaces to enable programmable and reversible binding^24^. This allows particles to be immobilized on solid substrates, assembled sequentially through inter-particle interactions, and subsequently released as defined multimeric products. Through this process, high-purity, low-order nanoparticle multimers have been generated, with dimers and trimers accounting for ∼70% of the total particle population^24^.

Building on nanomaterials assembly strategies, we establish a scalable workflow for controlled linking of AAV capsids. AAV capsids functionalized with single-stranded oligonucleotide linkers undergo stepwise coupling on solid substrates and, under optimized conditions, yield a narrow product distribution with dimers and trimers comprising ∼70% of the population. These linked AAVs enable coordinated co-delivery of genetic cargos, improving co-transduction efficiency and reducing cell-to-cell expression heterogeneity relative to dual-AAV delivery. This coordinated delivery supports reconstitution of large functional systems, exemplified by split prime editor 6d (PE6d) components with enhanced prime editing efficiency in primary neurons and in the mouse brain.

## Results

### Design of a workflow for controlled formation of linked AAVs

A workflow for controlled linking of AAV capsids was adapted from a previous report demonstrating stepwise assembly of oligonucleotide-functionalized nanoparticles on solid substrates^24^, with the introduction of an additional purification step to improve product homogeneity and enable recycling. In this workflow, two AAV vectors carrying distinct transgenes or split genetic cargos are defined as the linking targets and designated as Layer 1 AAV (L1-AAV) and Layer 2 AAV (L2-AAV).

In the first step, L1-AAVs, whose capsid surfaces are functionalized with two single-stranded oligonucleotides (A and B, each containing a 15 nucleotide (nt) main hybridization region and a 15 nt spacer), are immobilized onto streptavidin-coated magnetic bead substrates. The bead surfaces are functionalized with oligonucleotide A′, which contains a 15 nt sequence complementary to the hybridization region of oligonucleotide A and a 14 nt spacer terminated with biotin. Hybridization between the complementary regions of A and A′ enables immobilization of L1-AAVs on the substrate, and the bead-bound L1-AAVs are separated using a magnetic field (**Figure 1a**).

**Figure 1.**
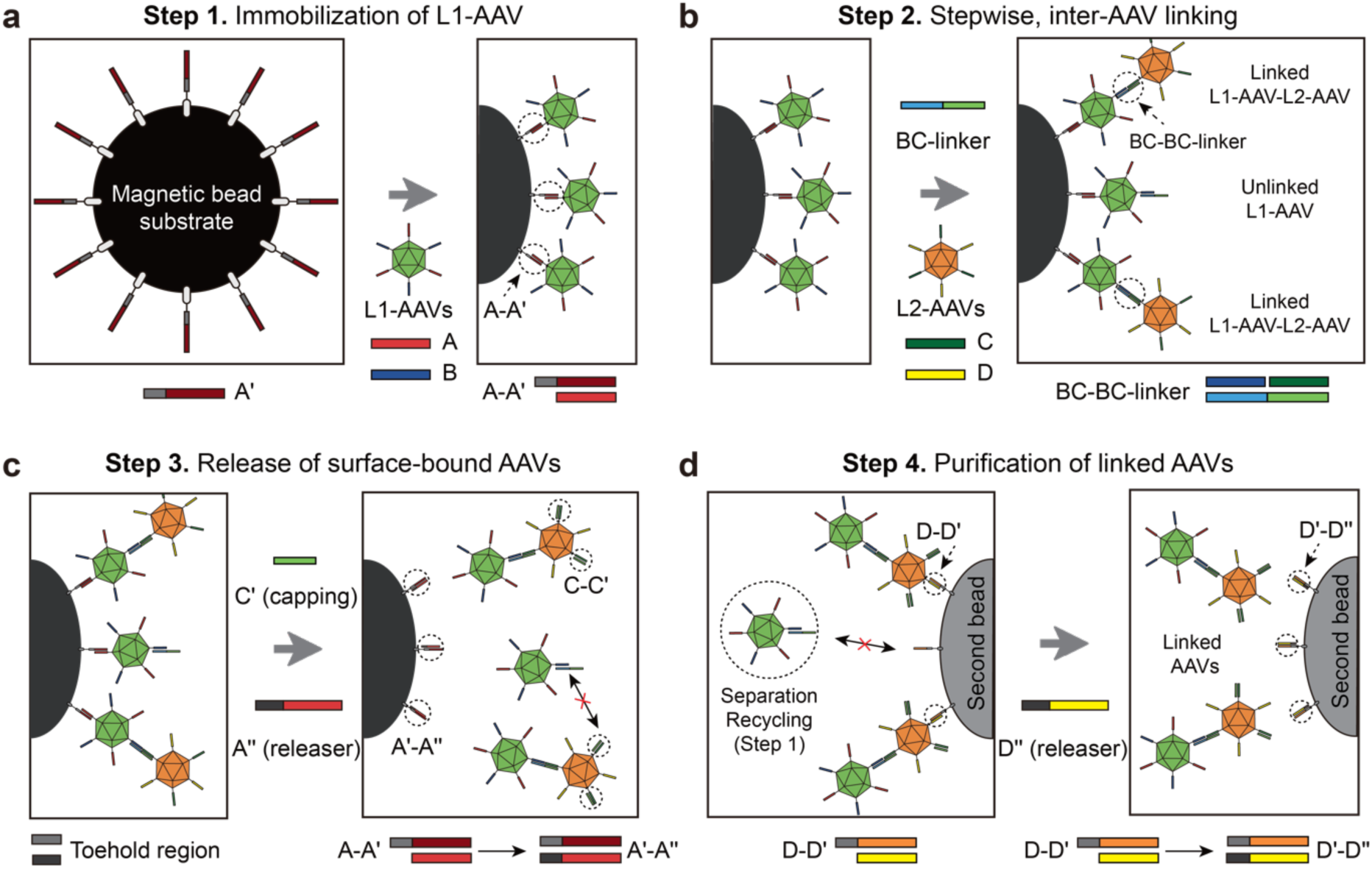
Workflow for linking AAV capsids and purification of linked AAVs. **(a)** L1-AAV anchoring: Oligonucleotide-modified L1-AAVs are immobilized on magnetic bead substrates through hybridization between capsid-displayed strand A and bead-anchored complementary strand A′. **(b)** Inter-AAV linking: L2-AAVs are introduced and linked to bead-bound L1-AAVs through the addition of a BC-linker. This results in linked AAVs together with residual unlinked L1-AAVs on the bead surface. Here, linked AAVs are illustrated as dimers for simplicity, although trimers and higher-order linked structures (n ≥ 4) may also form. **(c)** Release: Exposed linker sites on linked AAVs are passivated via a capping strand (C′) to prevent post-release re-association. Strand displacement mediated by a releaser strand (A′′) liberates AAVs from the bead surface. **(d)** Purification: The released AAVs are subjected to a second bead surface functionalized with D′ to enrich linked AAVs containing L2-AAV. A strand-displacement reaction mediated by D′′ yields purified linked AAVs, whereas unlinked L1-AAVs can be recycled.

Following immobilization, L1-AAVs are linked with L2-AAVs, whose capsid surfaces are functionalized with a second pair of oligonucleotides (C and D, each containing a 15-18 nt main hybridization region and a 12 nt spacer). Linking is initiated by the addition of BC-linker strands (32-53 nt) that are complementary to oligonucleotides B and C tethered to the respective AAV capsid surfaces. These linkers allow for the coupling of L2-AAVs onto substrate-bound L1-AAVs (**Figure 1b**). The surface-bound AAVs, consisting of linked AAVs together with residual unlinked L1-AAVs, are separated using a magnetic field, while unlinked L2-AAVs in the supernatant are collected for recycling.

After inter-capsid linking, residual oligonucleotide C strands on the linked AAVs are passivated by hybridization with a complementary capping strand (C′, 20 nt). After passivation, the surface-bound AAVs are released from the bead surface by the addition of a releaser strand (A′′, 23 nt). The A′′ strand contains an additional 8 nt relative to oligonucleotide A, which serves as a toehold region for strand displacement^27^. Through toehold-mediated strand displacement, a more thermodynamically stable duplex (A′ − A′′, 23 base pair (bp)) is formed compared to the original A − A′ duplex (15 bp), resulting in the release of the surface-bound AAVs from the bead substrates (**Figure 1c**).

Finally, the released AAVs are incubated with a second set of streptavidin-coated magnetic bead substrates functionalized with oligonucleotide D′, which contains a 15 nt sequence complementary to the hybridization region of oligonucleotide D and a 15 nt spacer terminated with biotin. Hybridization between D and D′enables the selective capture of linked AAVs containing L2-AAVs, while unlinked L1-AAV monomers remain in solution for recycling. After magnetic separation, the bead-bound AAVs are released by the addition of a releaser strand (D′′, 23 nt), which displaces the D − D′ duplex via toehold-mediated strand displacement in a manner analogous to the A′′-mediated release step (**Figure 1d**).

### Workflow optimization and fabrication of linked AAVs

To implement this workflow, we first selected two AAV9 vectors encoding distinct fluorescent reporters as model components, AAV9-CAG::mNeonGreen (L1-AAV) and AAV9-CAG::tdTomato (L2-AAV). The capsid surfaces of both L1-AAV and L2-AAV were functionalized with oligonucleotide linkers using inverse electron-demand Diels-Alder (iEDDA) click chemistry between tetrazine (Tz) and trans-cyclooctene (TCO). iEDDA click chemistry was selected over other bioorthogonal reactions because of its exceptionally fast kinetics, which can support efficient conjugation even at nanomolar reactant concentrations^28–30^. Briefly, AAV capsids were modified with Tz-PEG5-N-hydroxysuccinimide ester (Tz-PEG5-NHS) to target surface-exposed lysine residues, followed by conjugation with TCO-terminated oligonucleotides^31, 32^ (**Figure 2a**).

**Figure 2.**
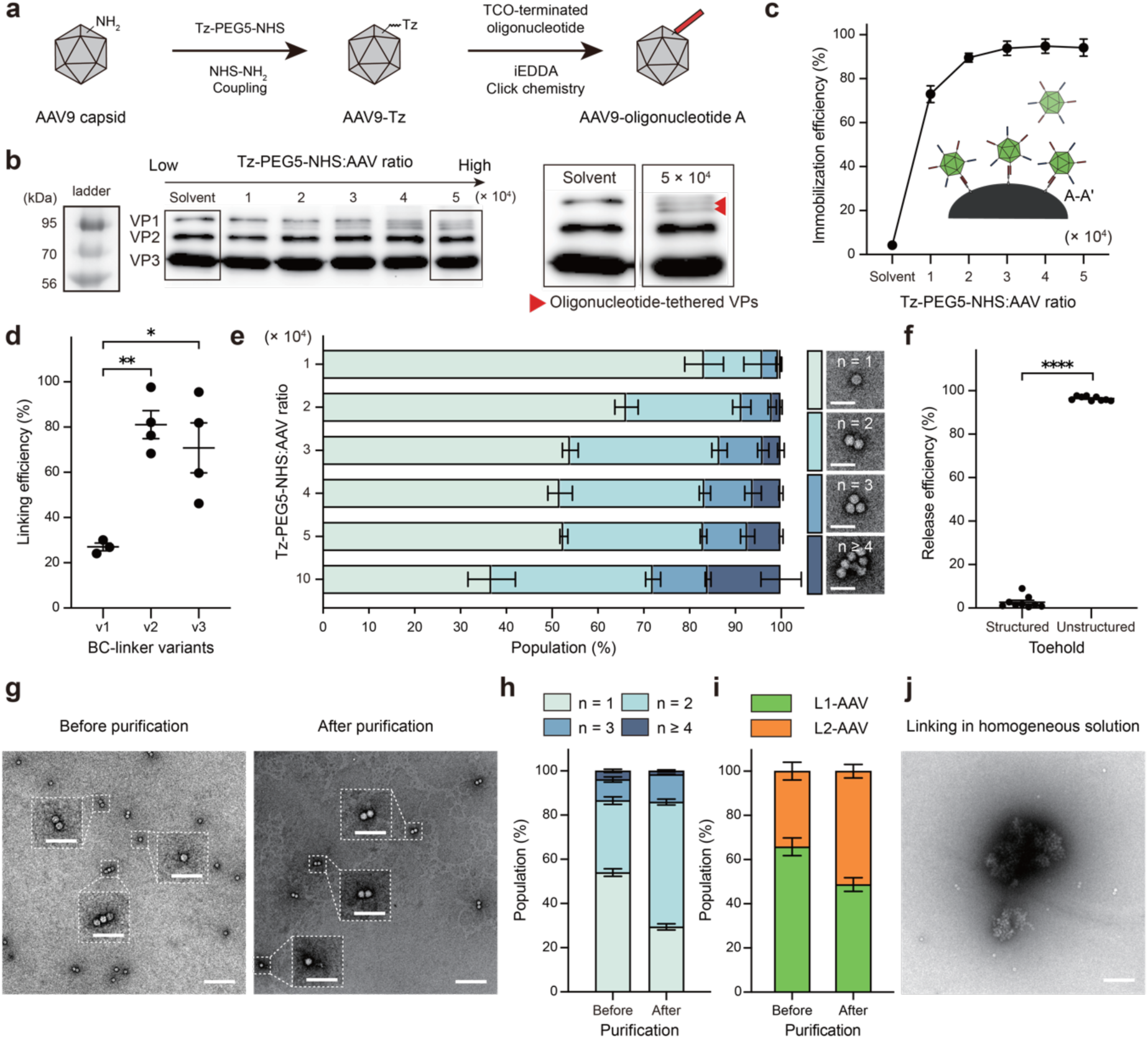
Optimization of the multi-AAV linking process. **(a)** Schematic illustration showing surface functionalization of AAV9 capsids via Tz-PEG5-NHS modification followed by iEDDA coupling with TCO-terminated oligonucleotides. **(b)** Western blot analysis showing progressive formation of higher-molecular-weight VP bands with increasing Tz-PEG5-NHS:AAV input ratios. Oligonucleotide-tethered capsid proteins are marked by red arrowheads. **(c)** L1-AAV immobilization efficiency on magnetic beads as a function of Tz-PEG5-NHS:AAV ratio. **(d)** Inter-AAV linking efficiency for different BC-linker variants. **(e)** TEM characterization of linking outcomes across varying oligonucleotide surface densities (left). Representative TEM images of AAV monomer, dimer, trimer, and higher-order aggregates (scale bars, 50 nm) (right). **(f)** Release efficiency of linked AAVs using releaser strands with structured or unstructured toeholds. **(g)** Representative TEM images of linked AAVs before and after purification (scale bars, 200 nm). Insets show magnified views of AAV monomers, dimers, or trimers (scale bars, 100 nm). **(h)** Quantitative population analysis of monomers, dimers, trimers, and higher-order linked AAVs before and after purification based on TEM imaging. **(i)** Genome composition analysis of linked AAVs before and after purification. **(j)** Representative TEM image of AAVs following the homogeneous linking process (scale bar, 200 nm). For Fig. 2c-f and 2h-i, data represent mean ± standard error of mean (s.e.m.) from at least three independent experiments. Statistical analysis for Fig. 2d was performed using one-way ANOVA. Statistical analysis for Fig. 2f was performed using an unpaired two-tailed Student’s t-test. Statistical significance is indicated as follows: \**p* < 0.05; \*\**p* < 0.01; \*\*\*\**p* < 0.0001.

Following conjugation, western blot analysis revealed new viral protein (VP) bands that were absent in unmodified AAVs. These newly appeared bands were shifted by ∼20 kDa relative to the main VP3 band, falling within the molecular weight range expected upon oligonucleotide conjugation. The intensity of the shifted bands increased as the Tz-PEG5-NHS:AAV ratio was raised from 1 × 10^4^ to 5 × 10^4^, with the TCO-oligonucleotide:AAV ratio fixed at 4 × 10^3^ (**Figure 2b**). Note that these ratios represent nominal input conditions used during the conjugation reaction and do not correspond to the absolute number of Tz-PEG5-NHS molecules or oligonucleotides attached per capsid^32, 33^.

To obtain functional evidence for oligonucleotide tethering on the AAV capsid surface, we next immobilized L1-AAVs on magnetic bead substrates functionalized with oligonucleotide A′. Immobilization efficiency was quantified by quantitative polymerase chain reaction (qPCR) analysis of mNeonGreen viral genomes (vg) retained on the bead surface. While control AAVs without oligonucleotide conjugation exhibited negligible binding to the bead substrate, L1-AAVs showed efficient immobilization, with efficiency strongly dependent on the Tz-PEG5-NHS:AAV ratio. Immobilization efficiency increased from 73.0 ± 3.8% at a ratio of 1 × 10^4^ and reached a plateau at ratios ≥3 × 10^4^, achieving 93.8 ± 3.2% retention after 1 h of incubation (**Figure 2c**). These results confirm that the conjugation chemistry effectively tethers oligonucleotides to the AAV capsid surface at levels sufficient to support substrate immobilization.

With L1-AAVs immobilized on magnetic bead substrates, we next linked L2-AAVs onto substrate-bound L1-AAVs. We hypothesized that inter-AAV linking efficiency would be strongly dependent on the design of the BC-linker, which bridges oligonucleotides B and C tethered on the L1-AAV and L2-AAV capsid surfaces, respectively. Based on this hypothesis, we first designed an initial BC-linker (BC-linker v1) using NUPACK predictions to achieve near-complete hybridization^34^. However, when tested with L1-AAVs and L2-AAVs functionalized at a Tz-PEG5-NHS:AAV ratio of 3 × 10^4^, BC-linker v1 resulted in limited inter-AAV linking. After 3 h of linking, qPCR analysis showed that only 27.0 ± 1.7% of input L2-AAV genomes were detected in the L1-AAV-bound fraction (**Figure 2d**). This limited linking efficiency likely reflects steric constraints and restricted conformational freedom of surface-tethered oligonucleotides, which are not captured in NUPACK solution predictions^35^.

To increase effective hybridization strength, we designed two extended BC-linker variants containing an additional 5-6 nt (BC-linker v2) or 10-11 nt (BC-linker v3) at both termini. With the same reaction time (3 h), inter-AAV linking efficiency increased to 81.1 ± 6.2% with BC-linker v2, consistent with enhanced hybridization strength. Further extension (BC-linker v3) slightly reduced inter-AAV linking efficiency to 70.8 ± 11.0% (**Figure 2d**), presumably due to increased steric interference between elongated linkers. These results highlight the importance of BC-linker design for efficient inter-AAV coupling, and BC-linker v2 was selected as the optimized linker for subsequent experiments.

Using the optimized BC-linker (v2), L1-AAVs and L2-AAVs functionalized at varying Tz-PEG5-NHS:AAV ratios were linked. The resulting constructs were subjected to the release process and subsequently analyzed by transmission electron microscopy (TEM). For quantitative TEM analyses, at least 500 released particles were analyzed from three independent experiments, and each product (monomers, dimers, trimers, and higher-order aggregates (n ≥ 4)) was counted as a single particle.

As the ratio increased from 1 × 10^4^ to 1 × 10^5^, the monomer fraction decreased from 83.2 ± 4.2% to 36.8 ± 5.2%, accompanied by a corresponding increase in linked species. Increasing the ratio from 1 × 10^4^ to 3 × 10^4^ led to progressive enrichment of low-order linked AAVs, with dimers and trimers reaching 32.6 ± 1.7% and 9.5 ± 1.2%, respectively. However, further increases beyond 3 × 10^4^ did not enhance dimer or trimer formation but instead promoted higher-order aggregation, with aggregate fractions rising from 3.9 ± 0.7% to 15.9 ± 4.4% (**Figure 2e**). The average number of capsids per aggregate also increased at higher Tz-PEG5-NHS:AAV ratios **(Supplementary Figs. 1-3)**, indicating enhanced multivalent crosslinking at elevated oligonucleotide densities. Based on these observations, a Tz-PEG5-NHS:AAV ratio of 3 × 10^4^ was selected as the optimal condition for generating low-order linked AAVs.

During this process, we identified an additional key parameter governing the efficiency of the release of substrate-bound AAVs. Specifically, the secondary structure of the toehold region in releaser A′′ strongly influenced the strand displacement efficiency. Releasers with predicted secondary structures in the toehold region exhibited markedly reduced recovery, with only 2.6 ± 0.9% of linked AAVs released after 3 h. In contrast, releasers designed to maintain unstructured toeholds enabled efficient strand displacement, resulting in 96.4 ± 0.3% recovery within 3 h (**Figure 2f**). These trends were consistent across multiple releaser designs (**Supplementary** Fig. 4), indicating that the releaser secondary structure is a critical determinant of the release efficiency. These results are aligned with a previous report showing that an unstructured releaser can effectively enhance toehold-mediated strand displacement efficiency^36^.

Released products generated under optimized conditions (BC-linker v2, Tz-PEG5-NHS:AAV ratio of ∼3 × 10^4^, and unstructured toehold releasers) were subsequently subjected to a purification step. TEM analysis demonstrated that purification reduced the monomer population from 54.0 ± 1.7% to 29.4 ± 1.4%, while increasing dimers and trimers from 32.6 ± 1.7% and 9.5 ± 1.2% to 56.5 ± 1.3% and 12.4 ± 0.1%, respectively, resulting in ∼70% low-order linked AAVs in the final population (**Figures 2g,h, and Supplementary Fig. 5**). Importantly, qPCR analysis showed that purification not only altered structural distribution but also restored compositional balance. Whereas released products prior to purification exhibited an excess of L1-AAV genomes, purification yielded near-equal genome copy numbers of L1-AAV and L2-AAV (**Figure 2i**).

Together, through optimization of oligonucleotide density, linker length, and releaser secondary structure, we established a workflow that generates low-order linked AAVs comprising ∼70% of the final population. These results contrast with linking reactions performed in homogeneous solution (Tz-PEG5-NHS:AAV ∼3 × 10^4^, BC-linker v2), which predominantly generated large AAV aggregates (**Figure 2j**). Notably, all linking, release, and purification steps were completed within 24 h using commercially available reagents **(Supplementary Fig. 6)**, and ≥93.0% of unlinked AAVs could be recycled for subsequent linking cycles **(Supplementary Fig. 7)**. Collectively, these features highlight the scalability and practical feasibility of the workflow for downstream biological applications.

### Coordinated delivery of AAV vectors through controlled linking

We next investigated whether our AAV linking strategy could enhance the coordinated delivery of AAV vectors in vitro. Using the previously defined mNeonGreen-encoding L1-AAV and tdTomato-encoding L2-AAV, linked AAVs were generated under optimized conditions (BC-linker v2, Tz-PEG5-NHS:AAV ratio of ∼3 × 10^4^, and unstructured toehold releasers), followed by purification. As controls, we included a physical mixture of the two unmodified AAV vectors, referred to as dual AAV, as well as a mixture of oligonucleotide-functionalized AAV vectors prepared at the same Tz-PEG5-NHS:AAV ratio but without the linking step, referred to as dual oligo-AAV (**Figure 3a**). The latter control was included to evaluate the potential effects of capsid surface modification on transduction behavior.

**Figure 3.**
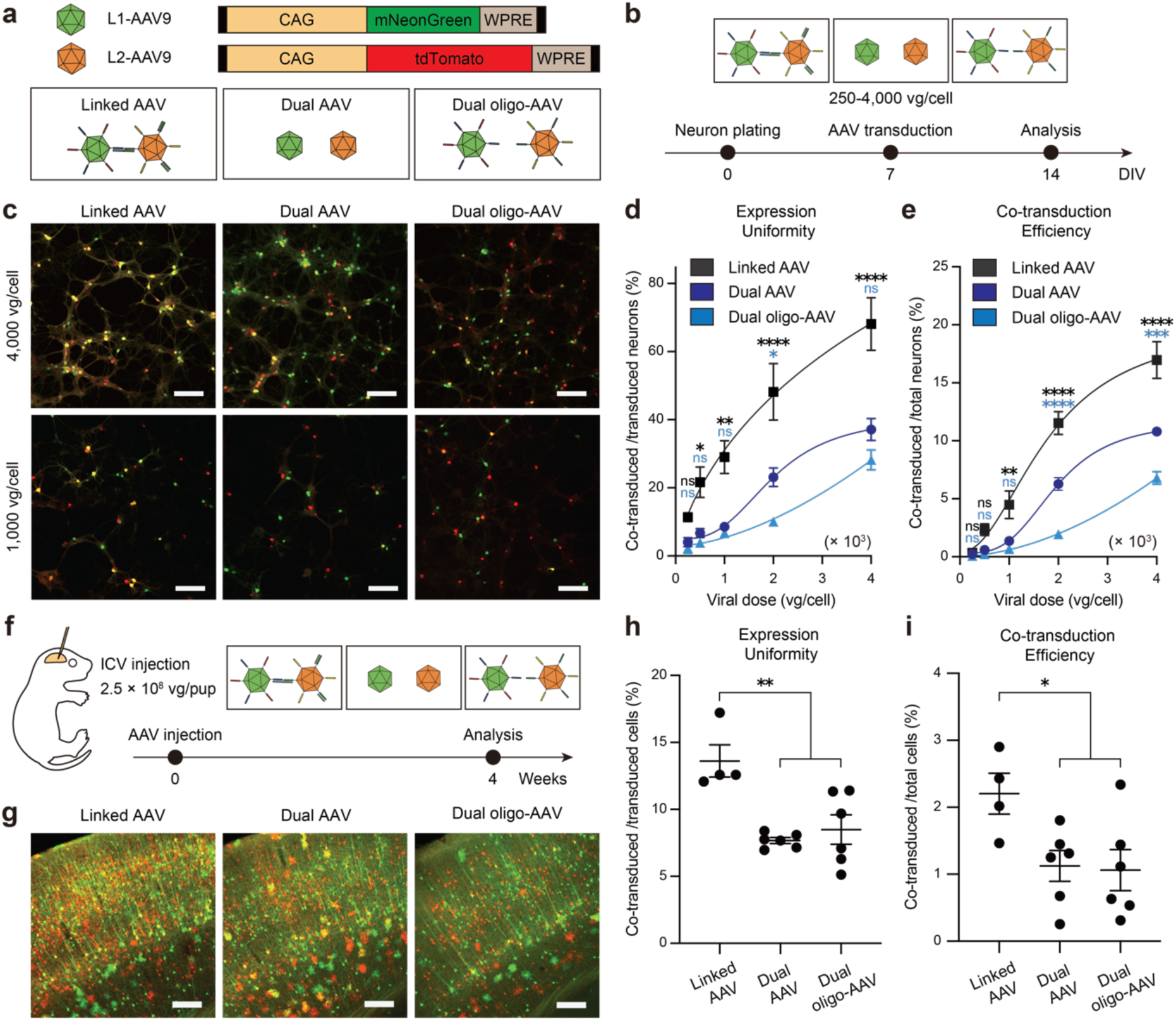
Linking-enhanced multi-AAV delivery in vitro and in vivo. **(a)** Illustration of linked AAVs and control groups, including dual AAV and dual oligo-AAV vectors encoding mNeonGreen (L1-AAV) and tdTomato (L2-AAV). **(b)** Experimental timeline for in vitro neuronal transduction at various viral doses. **(c)** Representative fluorescence images of primary hippocampal neurons transduced with linked AAV, dual AAV, or dual oligo-AAV at 4,000 (top) and 1,000 (bottom) vg/cell (scale bars, 200 µm). **(d)** Quantification of expression uniformity across viral doses, defined as the proportion of neurons co-expressing mNeonGreen and tdTomato among total transduced neurons within defined ROIs. **(e)** Quantification of co-transduction efficiency across viral doses, defined as the fraction of neurons co-expressing both reporters among total neurons within each ROI. **(f)** In vivo experimental scheme for neonatal mouse ICV injection of linked AAVs and control vectors, followed by analysis four weeks post-injection. **(g)** Representative fluorescence images of neocortical ROIs following ICV delivery of linked AAVs or control vectors (scale bars, 200 µm). **(h)** Quantification of expression uniformity in neocortical ROIs four weeks post-injection, showing increased proportions of co-transduced cells among total transduced cells in the linked AAV group compared with controls. **(i)** Quantification of co-transduction efficiency in neocortical ROIs, confirming preservation of enhanced coordinated delivery in vivo with linked AAVs. For Fig. 3d-e and 3h-i, results represent mean ± s.e.m. derived from at least three independent biological replicates. For Fig. 3d-e, statistical significance was performed using two-way ANOVA followed by Dunnett’s multiple-comparison test, comparing each group with dual AAV control at each viral dose. For Fig. 3h-i, statistical analysis was performed using one-way ANOVA. Statistical significance is indicated as follows: *n.s.*, not significant; \**p* < 0.05; \*\**p* < 0.01; \*\*\**p* < 0.001; \*\*\*\**p* < 0.0001.

Primary hippocampal neurons were transduced with linked AAVs, dual AAVs, and dual oligo-AAVs across a range of viral doses from 250 to 4,000 vg/cell (**Figure 3b**). Expression uniformity and overall co-transduction efficiency were quantified based on the numbers of neurons expressing both mNeonGreen and tdTomato (co-transduced), neurons expressing only one reporter (partially transduced), and total neurons within defined regions of interest (ROIs). For each condition, at least 1,000 neurons were analyzed from 5 ROIs in each of three independent experiments.

Across all tested viral doses, linked AAVs showed significant enhancement of expression uniformity compared with both control groups (**Figure 3c and Supplementary Fig. 8**). At 4,000 vg/cell, linked AAVs achieved 68.1 ± 7.7% co-transduction among transduced neurons, corresponding to 1.8-fold and 2.4-fold improvements over dual AAV and dual oligo-AAV controls, respectively (**Figure 3d**). Consistent with this improvement, the fractions of mNeonGreen-only and tdTomato-only neurons among transduced neurons were 17 ± 1.7% and 14 ± 6.3% in the linked AAV group, corresponding to 0.5-fold of the levels observed in the dual AAV and dual oligo-AAV control groups (**Supplementary Fig. 9**). Linked AAVs also increased the proportion of co-transduced neurons among total neurons. At 4,000 vg/cell, 17.0 ± 1.6% of total neurons were co-transduced in the linked AAV group, compared with 10.8 ± 0.4% and 6.8 ± 0.6% in the dual AAV and dual oligo-AAV control groups, respectively (**Figure 3e**).

The enhancement in coordinated delivery by linked AAVs became more pronounced at lower viral doses. At 1,000 vg/cell, linked AAVs exhibited 3.4-fold and 4.3-fold higher expression uniformity compared with dual AAV and dual oligo-AAV controls, respectively (**Figure 3d**). At the same dose, the proportion of co-transduced neurons relative to total neurons increased by 3.3-fold and 6.6-fold, respectively (**Figure 3e**). These findings suggest that, at lower viral doses, the probability of simultaneous delivery of AAV vectors becomes increasingly constrained under conventional dual AAV conditions, whereas linked AAVs maintain a distinct level of coordinated delivery.

As a result of enhanced coordinated delivery, linked AAVs achieved comparable co-transduction at reduced viral doses relative to dual AAVs, while maintaining higher expression uniformity. For example, at 2,000 vg/cell, linked AAVs achieved co-transduction efficiency (11.5 ± 1.0%) comparable to that of dual AAVs (10.8 ± 0.4%) at 4,000 vg/cell (**Figure 3e**). Under the same condition, expression uniformity was higher in the linked AAV group (48.2 ± 8.3%) than in dual AAV controls (37.1 ± 3.2%) (**Figure 3d**).

To assess whether the coordinated delivery achieved with linked AAVs in vitro can be preserved in vivo, we performed unilateral intracerebroventricular (ICV) injections of linked AAVs and corresponding control vectors into neonatal mice. Given that lower viral doses produced more pronounced differences in coordinated delivery in vitro, we selected a total dose of 2.5 × 10^8^ vg/pup for neonatal ICV injection, which is substantially lower than the commonly used range of 10^10^-10^11^ vg/pup^37–39^ (**Figure 3f**).

Four weeks after injection, robust fluorescent signals were predominantly observed in the neocortex across all experimental groups (**Supplementary Fig. 10)**. Expression uniformity was calculated as the proportion of co-transduced cells among total transduced cells, and co-transduction efficiency was calculated as the fraction of co-transduced cells relative to total cells within defined ROIs of the neocortex (**Figure 3g**). Consistent with the in vitro findings, linked AAVs resulted in markedly higher expression uniformity (13.6 ± 1.2%), whereas dual AAV and dual oligo-AAV controls exhibited substantially lower uniformity (7.7 ± 0.2% and 8.5 ± 1.1%, respectively) (**Figure 3h**). In addition, co-transduction efficiency within the ROI reached 2.2 ± 0.3% in the linked AAV group, exceeding that observed in dual AAV and dual oligo-AAV controls (1.1 ± 0.2% and 1.1 ± 0.3%, respectively) (**Figure 3i**). Collectively, these results demonstrate that linked AAVs preserve enhanced coordinated delivery within a brain environment.

### Enhanced gene editing via linked AAVs in vitro and in the mouse brain

To demonstrate the functional feasibility of our AAV linking strategy, we applied it to deliver a split prime editor system that has previously been implemented using dual AAV approaches^10, 11^. In this established configuration, one AAV9 vector encodes the C-terminal fragment of PE6d fused to the C-terminal Npu split intein, while the second AAV9 vector encodes the complementary N-terminal intein fragment fused to the remaining portion of PE6d (**Figure 4a**). Coordinated delivery enables intein-mediated protein splicing, reconstituting a full-length PE6d capable of inserting a 42 bp loxP-containing sequence at the murine *Dnmt1* locus^11^ (**Figure 4b**).

**Figure 4.**
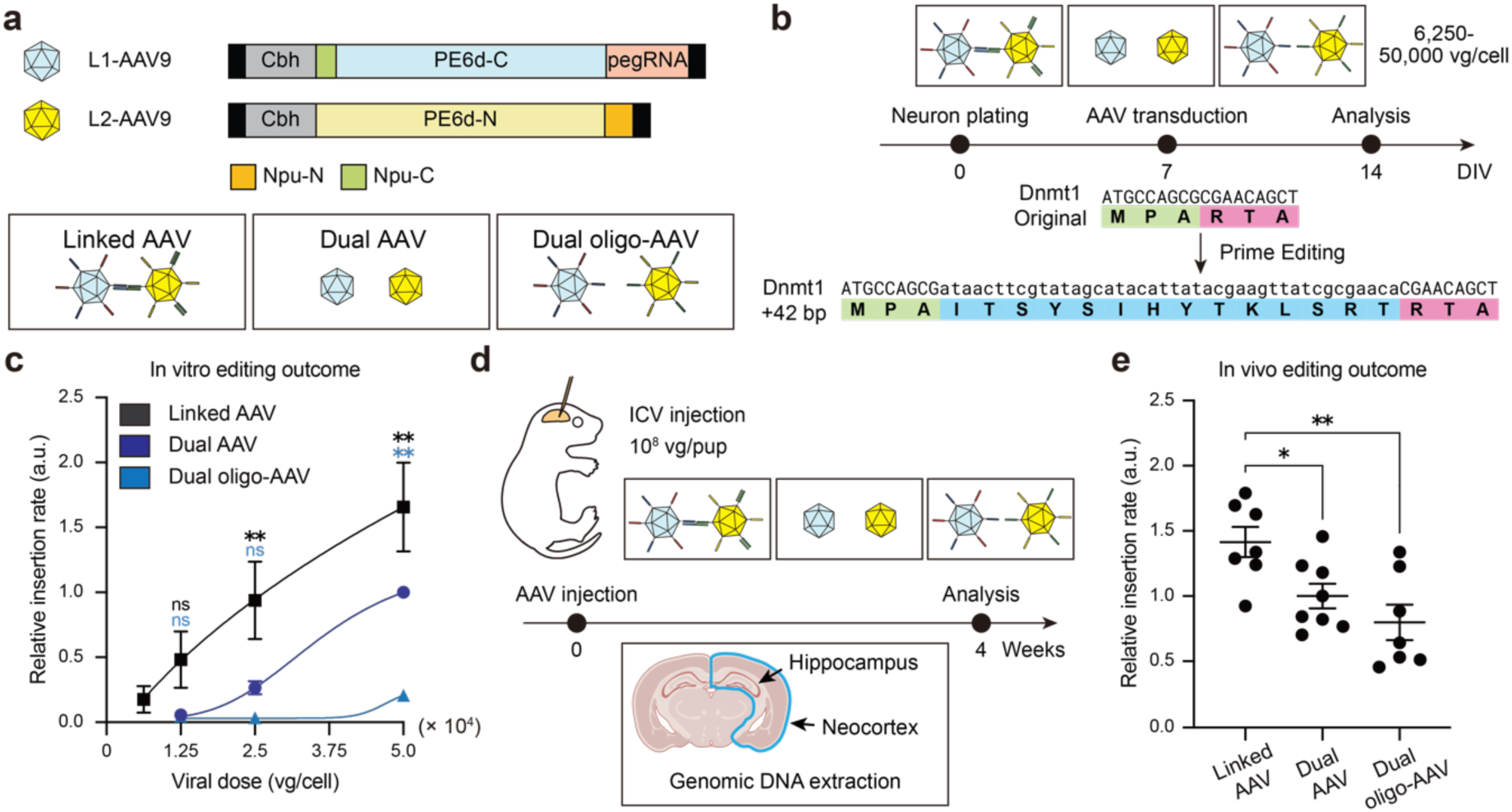
Improved prime editing via coordinated split gene delivery. **(a)** Schematic of linked AAV and control configurations (dual AAV and dual oligo-AAV) for delivering complementary fragments of split PE6d. **(b)** Experimental workflow for prime editing in primary hippocampal neurons following delivery of linked AAVs or control vectors. Coordinated dual-vector delivery induces intein-mediated reconstitution of functional PE6d, enabling targeted insertion of a 42 bp loxP-containing sequence at the *Dnmt1* locus. **(c)** Dose-dependent relative insertion level in primary hippocampal neurons transduced with linked AAVs or control vectors. Relative insertion abundance was quantified by qPCR and normalized within each experimental batch to the △Ct value obtained from dual AAV at 50,000 vg/cell. At 6,250 vg/cell, control groups did not yield reliable qPCR signals (Ct > 30) and were therefore excluded from the plot. **(d)** Schematic of the in vivo experimental design for neonatal ICV delivery of linked AAV and control split PE6d vectors, followed by genomic DNA isolation from cerebral cortex. The brain slice cartoon was created using BioRender. **(e)** Quantification of relative insertion abundance in vivo four weeks post-injection, demonstrating enhanced prime editing following linked AAV delivery compared with control groups. qPCR data were normalized to the mean value of the dual AAV group. For Fig. 4c and 4e, data are shown as mean ± s.e.m. from at least three independent biological replicates. For Fig. 4c, statistical significance was performed using two-way ANOVA followed by Dunnett’s multiple-comparison test, comparing each group with dual AAV control at each viral dose. For Fig. 4e, statistical analysis was performed using one-way ANOVA. Statistical significance is indicated as follows: *n.s.*, not significant; \**p* < 0.05; \*\**p* < 0.01.

Based on this foundation, AAV9 vectors encoding the split PE6d components were linked under optimized conditions and subsequently purified. Primary hippocampal neurons were then transduced with linked AAVs across a broad range of viral doses (6,250–50,000 vg per cell), with dual AAV and dual oligo-AAV groups serving as controls. Editing outcomes were assessed by measuring the relative abundance of the inserted 42 bp sequence at the *Dnmt1* locus using qPCR (**Figure 4b**). This analysis provides a comparative measure of editing output between groups rather than an absolute editing frequency.

Across the tested dose range, linked AAVs consistently produced higher insertion levels than control groups, with the magnitude of improvement becoming more pronounced as the viral input decreased. At the highest viral dose (50,000 vg/cell), linked AAVs showed a 1.7-fold increase over dual AAV controls, indicating a relatively modest benefit under saturating conditions. At 25,000 and 12,500 vg/cell, however, linked AAVs achieved 3.5-fold and 11.0-fold higher insertion signals relative to dual AAV controls, respectively. At 6,250 vg/cell, insertion signals in the control groups fell below the reliable detection limit of qPCR (cycle threshold (Ct) value > 30), whereas linked AAVs still yielded detectable levels of gene insertion. Notably, dual oligo-AAV controls consistently exhibited markedly lower insertion levels than both linked AAVs and dual AAV controls across all tested doses (**Figure 4c**), indicating that the enhanced editing performance arises from capsid linking rather than oligonucleotide conjugation.

To assess whether this functional advantage can be preserved in vivo, neonatal mice received ICV injections of linked or control split PE6d AAV vectors. Because the disparity in editing efficiency became more pronounced at lower viral inputs in vitro, we employed a total dose of 1 × 10^8^ vg/pup, which is substantially below the commonly used range (10^10^-10^11^ vg/pup)^10, 11^. Four weeks after administration, genomic DNA was isolated from the cerebral cortex in the injected hemisphere, where robust fluorescent signals were observed in the preceding experiment (**Supplementary Fig. 10**), and the relative abundance of the targeted 42 bp insertion was quantified and compared across groups (**Figure 4d**). Consistent with the primary neuron experiments, linked AAVs yielded statistically higher insertion signals in vivo (1.4-fold and 1.8-fold relative to dual AAV and dual oligo-AAV controls, respectively) (**Figure 4e**). These results demonstrate that split gene delivery mediated by AAV linking is feasible in vivo and supports effective reconstitution of a multi-component genome editing system.

Taken together, these results demonstrate that linked AAVs can improve coordinated multi-vector delivery, boosting not only reporter co-expression but also functional genome editing. By increasing the probability that complementary vectors reach the same cell, linked AAVs sustain measurable editing activity under conditions in which conventional dual AAV approaches are limited.

### Conclusion

In this work, we developed a strategy for the controlled linking of nanoscale viral capsids, drawing on principles from nanomaterials engineering^23–26^. Rather than relying on conventional solution-phase assembly, which often yields heterogeneous products, our approach confines the linking process to solid interfaces, enabling controlled multimer formation. Oligonucleotides serve as key linkers in this process, supporting precise, stepwise assembly of capsids on the surface, as well as efficient release and downstream purification. Under optimized conditions, this approach provides a scalable process that yields AAV dimers and trimers comprising ∼70% of the final population, with high recyclability (≥93.0%) and short processing times (∼24 h). By coupling viral particles prior to administration, AAV linking increases the probability of co-delivery into the same cell, thereby enhancing expression uniformity and co-transduction efficiency, and facilitating reconstitution of split genetic systems in vitro and in vivo.

For broader application of our AAV linking strategy, its compatibility with engineered AAV capsids designed for systemic delivery should be considered. In our pilot experiments with brain-tropism-enhanced variants (AAV.CAP-B10 and AAV9P31)^40, 41^ and adult mice, oligonucleotide modification markedly reduced the brain tropism of CAP-B10, whereas 9P31 relatively retained detectable brain transduction after capsid modification **(Supplementary Fig. 11)**. This difference likely reflects the sequence composition of the engineered motifs: the 9P31 sequence inserted for brain tropism (WPTSYDA) lacks lysine residues for oligonucleotide tethering, whereas the inserted CAP-B10 sequences (DGAATKN, TLAVPFK) contains lysines^42, 43^. The absence of surface-exposed lysine residues within engineered receptor interaction motifs may therefore provide a guideline for identifying capsids compatible with the linking strategy.

Our systemic delivery experiments with linked AAV9P31 vectors and adult mice showed improved expression uniformity among transduced cells, although overall brain transduction remained lower than that observed with unmodified vectors **(Supplementary Figs. 11 and 12)**. These findings indicate that further optimization will be required to preserve the systemic delivery efficiency of engineered capsids. This may be achieved through site-selective conjugation strategies, such as labeling through noncanonical amino acids, to minimize interference between oligonucleotide linkers and structural regions required for engineered tropism^44–46^.

In summary, we establish a framework for linking AAV capsids into defined multi-particle architectures, enabling more reliable co-delivery of split genetic cargos, as exemplified by prime editing applications. We anticipate that further refinement of capsid conjugation strategies and controlled assembly may expand the utility of this platform for multi-vector delivery and facilitate the development of AAV-based therapies requiring delivery of large genetic payloads. In addition, beyond the applications demonstrated here, our capsid linking strategy may provide a general framework for engineering higher-order viral assemblies for diverse delivery and functional applications.

## Methods

### Chemicals

The chemicals used in this study were as follows. Sodium chloride (ACS regent, 99.0%; NaCl, S9888), magnesium chloride hexahydrate (≥ 99%; MgCl_2_·6H_2_O, M9272), calcium chloride dihydrate (≥ 99%; CaCl_2_·2H_2_O, 223506), sodium carbonate monohydrate (≥99.5%; Na_2_CO_3_, S4132), N-lauroylsarcosine sodium salt (≥97.0%, HPLC; 61745), 5-fluoro-2’-deoxyuridine (FUDR; F003-100MG), paraformaldehyde powder (PFA, 95%; 158127), Tween 20 (Molecular Biology, viscous liquid; P9416), 2-mercaptoethanol (Molecular Biology, 99% (GC/titration); M3148), and glycerol (≥ 99%; G6279) were purchased from Sigma-Aldrich. Pluronic F-68 Non-ionic Surfactant (100×; 24040032) was purchased from Gibco. 0.5 M Tris-HCl (pH 6.8; ETB014-500), 1.5 M Tris-HCl (pH 8.8; ETB015-500), and 10% SDS (EBS006-500) were purchased from Enzynomics. 0.5 M EDTA solution (pH 8.0; ML005-01), PBS (pH7.4; ML008-01), and water (Pure WFI; ML019-02) were purchased from WELGENE. All oligonucleotides were synthesized by Bioneer unless otherwise specified, and were purified by HPLC. TCO-conjugated oligonucleotides were synthesized by Integrated DNA Technologies and purified by HPLC.

### Plasmid DNA

The following plasmids were used for AAV production: pAAV2/9n (Addgene plasmid #112865; http://n2t.net/addgene:112865; RRID: Addgene_112865) and pAdDeltaF6 (Addgene plasmid #112867; http://n2t.net/addgene:112867; RRID: Addgene_112867). These plasmids were gifts from James M. Wilson. pAAV-CAG-tdTomato (codon diversified) was a gift from Edward Boyden (Addgene plasmid #59462; http://n2t.net/addgene:59462; RRID: Addgene_59462). pAAV-CAG-2xNLS-tdTomato-WPRE-hGHpolyA was a gift from Guoping Feng (Addgene plasmid #192552; http://n2t.net/addgene:192552; RRID:Addgene_192552). A v3em-Cterm-PE6d-dualU6-Dnmt1-PE3-loxp (Addgene plasmid #207863; http://n2t.net/addgene:207863; RRID: Addgene_207863) and v3em-Nterm-PE2max (Addgene plasmid #198734; http://n2t.net/addgene:198734; RRID: Addgene_198734) were gifts from David Liu^10, 11^.

Standard molecular cloning techniques were used to generate DNA constructs in this study. For pAAV-CAG-mNeonGreen, the backbone of pAAV-CAG-tdTomato was digested with BamHI and HindIII. The insert fragment encoding mNeonGreen was amplified using Phusion High-Fidelity DNA Polymerase (NEB, M0530L) with pAAV-hSyn1-mNeonGreen (Addgene plasmid #99135; http://n2t.net/addgene:99135; RRID: Addgene_99135) as a template and ligated using T4 DNA ligase (NEB, M0202S). For pAAV-CAG-mNeonGreen-NLS, the backbone of pAAV-CAG-mNeonGreen was digested with EcoRI. The insert fragment encoding the NLS motif was amplified using Phusion High-Fidelity DNA Polymerase with pAAV-CAG-2xNLS-tdTomato-WPRE-hGHpolyA. Insert fragment and backbone were ligated using NEBuilder HiFi DNA assembly master mix (NEB, E2621L). pAAV-9P31 was generated using the pAAV2/9n backbone with the Q5 Site-Directed Mutagenesis Kit (NEB, E0554S) and primers. The primer sequences used for cloning and mutagenesis are provided in Supplementary Table 1. All constructs were verified by Sanger sequencing.

### AAV production

AAVs were produced and purified according to a published method^47^ with minor modifications. In brief, HEK293T cells were co-transfected with a transfer plasmid, a packaging plasmid and pAdDeltaF6 as a helper plasmid with PEI Max (MW 40,000; Polyscience, 24765-100). Supernatants were collected at 72 and 120 h post-transfection, and transfected cells were harvested at 120 h. Viral particles from supernatants were concentrated by poly(ethylene glycol) (BioUltra, 8000; Sigma-Aldrich, 89510-250G-F) precipitation. Cell pellets were lysed, and residual DNA was removed using Salt Active Nuclease (SAN; New England Biolabs, M0764S) and SAN digestion buffer (500 mM NaCl, 40 mM Tris base, and 10 mM MgCl_2_ in water). The lysate was clarified and combined with concentrated supernatant fractions. AAVs were purified using an iodixanol (Optiprep; Serumwerk, AXS-1114542) gradient from 15% to 60% (wt/vol) followed by ultracentrifugation with a Type 70 Ti-fixed angle titanium rotor (Beckman Coulter, 337922) at 350,000 × g for 2.4 h. The 40% fraction and the 40/60% interface were collected, then buffer-exchanged and concentrated using a 100 kDa Amicon Ultra filter unit (Sigma-Aldrich, UFC910008). Purified AAVs were stored at 4 °C for up to 6 months.

### AAV titration

AAV vector genome titers were quantified according to a published method^47^. Briefly, 2 µl of AAV sample was incubated with 100 µl of DNase I mixture (5 U of DNase I (Roche, 4716728001), 2 mM CaCl_2_, 10 mM Tris-HCl, and 10 mM MgCl_2_ in water) at 37 °C for 1 h to degrade unpackaged DNA. DNase I was inactivated by adding 5 µl of 0.5 M EDTA, followed by incubation at 70 °C for 10 min. Subsequently, 120 µl of proteinase K solution (120 µg of proteinase K (Roche, 3115887001), 1 M NaCl and 1% (wt/vol) N-lauroylsarcosine sodium salt in water) was added, and samples were incubated at 50 °C for 2 h to digest viral capsid proteins. Proteinase K was inactivated by heating at 95 °C for 10 min. Samples were then cooled at 4 °C for 5 min and diluted by mixing 3 µl of each sample with 897 µl of water. qPCR was performed using THUNDERBIRD Next SYBR qPCR mix (Toyobo, TOQPX-201) with appropriate standard samples. Vector genome titers were calculated based on a standard curve generated from serial dilutions of digested plasmid DNA. Primer sequences for each AAV titration are provided in Supplementary Table 1.

### Surface modification of AAV capsids with oligonucleotides

To conjugate oligonucleotides (A, B, C, and D) onto the AAV capsid surface, 2.5 µl Tz-PEG5-NHS (Sigma-Aldrich, 900913) dissolved in anhydrous DMSO was added to 1 nM AAV sample in NHS reaction buffer (PBS, pH adjusted to 8.3 with 0.1 M Na_2_CO_3_ solution). The total reaction volume was adjusted to 250 µl. After 1 h incubation at room temperature, an additional 2.5 µl of Tz-PEG5-NHS was added, followed by further incubation for 1 h. Following Tz modification, samples were buffer-exchanged into AAV storage buffer (0.001% Pluronic F-68 in PBS) using an Amicon Ultra centrifugal filter unit (100 kDa MWCO; Millipore, UFC10096). Buffer exchange was performed five times at 3,000 × g for 15 min each. For oligonucleotide conjugation, 20 µl of TCO-oligonucleotide mixture (1:1 ratio of 100 µM oligonucleotide A (or C) and 100 µM oligonucleotide B (or D)) was then added to Tz-modified AAVs. The total reaction volume was adjusted to 250 µl in AAV storage buffer, and samples were incubated for 6 h at room temperature with gentle rotation at 15 rpm (Multi Bio RS-24 Programmable rotator; Biosan). Excess, unbound oligonucleotides were removed by five washing steps using an Amicon Ultra centrifugal filter unit (100 kDa MWCO) with the AAV linking buffer (163 mM NaCl, 12.5 mM MgCl_2_, 0.001% Pluronic F-68 in PBS).

### Western blot

For western blot analysis, AAV samples equivalent to 1 × 10^9^ vg per lane, as determined by AAV genome titration, were loaded. Samples were mixed with 6× Laemmli sample buffer (375 mM Tris-HCl, 9% SDS, 50% glycerol, 9% 2-mercaptoethanol, and 0.03% bromophenol blue at pH 6.8) and heated at 95 °C for 5 min. Samples were then run on a 12% SDS-polyacrylamide gel with running buffer (25 mM Tris, 190 mM glycine, and 0.1% SDS in water) at 100 V for 85 min and transferred onto a 0.45 µm PVDF membrane (Amersham Hybond P 0.45 PVDF; Whatman, 10600023) using transfer buffer (25 mM Tris, 190 mM glycine, and 20% methanol in water) at 100 V for 85 min. Membranes were blocked for 1 h at room temperature in blocking buffer (5% (wt/vol) nonfat dry milk (Bio-Rad, BR1806404), 20 mM Tris, 150 mM NaCl, and 0.1% Tween 20 at pH 7.5). Membranes were next incubated with anti-AAV B1 antibody (1:250; PROGEN, 690058S) for 1 h at room temperature, followed by HRP-conjugated goat anti-mouse antibody (1:10,000; Abcam, ab205719) for 1 h. Membranes were washed three times with TBST between incubations. Protein bands were finally detected using TOPview ECL Femto Western Substrate (Enzynomics, EOE003S) and imaged with a ChemiDoc Imaging System (Bio-Rad, 12003153).

### Oligonucleotide tethering to magnetic bead substrates

Dynabeads MyOne Streptavidin C1 magnetic beads (Invitrogen, 65001) were used to immobilize biotinylated oligonucleotides. First, 50 µl of magnetic beads were washed three times with Binding and Washing (B&W) buffer (5 mM Tris-HCl, 500 µM EDTA, and 1 M NaCl at pH 7.5). The washed beads were resuspended in 100 µl of 2× B&W buffer and mixed with 100 µl of biotin-terminated oligonucleotides (*A*′ or *D*′) (100 µM). The mixture was incubated at room temperature for 1 h at 15 rpm for oligonucleotide immobilization onto the magnetic bead substrates. After immobilization, beads were collected using a magnetic rack for 5 min and washed twice with B&W buffer, twice with EDTA-pluronic buffer (0.1 mM EDTA and 0.001% Pluronic F-68 in PBS), and twice with AAV linking buffer. Finally, beads were resuspended in 50 µl of AAV linking buffer and stored at 4 °C for up to 2 weeks.

### Fabrication of linked AAVs

For multi-AAV linking, all the reactions were performed in Protein LoBind tubes (2.0 ml, PCR clean; Eppendorf, 30108132) using AAV linking buffer at room temperature. During incubation, samples were continuously rotated at 15 rpm using a Multi Bio RS-24 programmable rotator. The total reaction volume was 250 µl unless otherwise specified.

For L1-AAV immobilization, 0.25 nM L1-AAVs were incubated with 125 µg (12.5 µl) of *A*′ functionalized magnetic beads for 1 h. After magnetic separation and removal of the supernatant, 2.5 µl of 10 µM BC-linker was then added and incubated for 1 h. Following the three washing steps with AAV linking buffer, 0.25 nM L2-AAVs were added and incubated for 3 h for inter-AAV linking. Subsequently, 2.5 µl of 10 µM *C*′ was added and incubated for 1 h. After three additional washing steps, 9.38 µl of 100 µM *A*″ was added and incubated for 3 h to release the AAVs bound to the magnetic bead substrate.

For purification, the supernatant collected after *A*″ oligonucleotide addition was incubated with 125 µg (12.5 µl) of *D*′functionalized magnetic beads for 1 h. After three washing steps, 12.5 µl of 100 µM *D*″ was added and incubated for 1 h to release the AAVs bound to the magnetic bead substrate. Finally, linked AAVs were collected from the supernatant and stored at -80 °C until further use.

### TEM analysis of linked AAVs

Linked AAVs were deposited onto formvar/carbon supported copper grids (Electron Microscopy Sciences; FCF-200-Cu). Grids were glow-discharged for 30 s at 25 mA using a PELCO easiGlow Glow Discharge Cleaning System (TED PELLA) prior to sample application. 5 µl of AAV sample was loaded onto each grid and incubated for 1 min. Excess liquid was blotted with filter paper, and grids were washed with two drops of water. Negative staining was performed with two drops of 2% (wt/vol) uranyl acetate solution for 1 min. After removal of excess stain, the stained grids were air-dried for 5 min. Images were acquired using a Talos F200X G2 transmission electron microscope (Thermo Scientific) operated at 200 kV at the KAIST Analysis Center for Research Advancement. Images were recorded at a nominal magnification of 11,000×.

### Animal care

All animal experiments were conducted in accordance with the guidelines of the Institutional Animal Care and Use Committee (IACUC) at KAIST under approved protocols KA2024-147-v1 and KA2025-050-v1. Pregnant female C57BL/6N and female ICR mice were obtained from KOATECH for neonatal ICV injections and primary neuron cultures. Six-week-old male C57BL/6N mice were purchased from KOATECH and used for retro-orbital (RO) injections. Mice were housed under a 12 h light/dark cycle at 20-25 °C and 30-70% relative humidity with ad libitum access to standard rodent diet and water.

### Primary neuron culture

Primary hippocampal neurons were prepared based on a published method^48^, with minor modifications. Postnatal day 1 C57BL/6N pups were anesthetized by hypothermia on ice and euthanized by cervical dislocation after confirming the absence of reflexes. Brains were rapidly removed and placed in cold dissection buffer (5.5 mM glucose in PBS) at 4 °C. Hippocampi were isolated and incubated in 10 ml of papain digestion solution (10 mg BSA, 10 mM glucose, 100 µg DNase I (Roche, 11284932001), and 5 mg papain (Papain from Carica papaya; Sigma-Aldrich, 76216-250MG) in 10 ml PBS sterilized using a 0.22 µm syringe filter). Tissues were incubated at 37 °C for 15 min with gentle inversion every 5 min. After enzymatic digestion, papain solution was removed and tissues were washed twice with 5 ml of plating medium (0.02% B27 supplement (Gibco, 17504044), 0.1% Fetal bovine serum (WELGENE, S001-01), 2.5 µg/ml amphotericin B (Sigma-Aldrich, A2942-100ML), 100 µg/ml gentamicin (Sigma-Aldrich, G1272-10ML) in neurobasal A medium (Thermofisher, 10888022)). Tissues were resuspended in 2 ml of plating medium and mechanically dissociated using a fire-polished Pasteur pipette (HILGENBERG, HG.3150101) until a homogeneous cell suspension was obtained. Primary hippocampal neurons were counted using trypan blue and seeded at a density of 150,000 cells per well in poly-D-lysine-coated 24-well culture plates. Cultures were maintained with culture medium (1 ml of B27 supplement, 150 µl of 100× Glutamax (Gibco; 35050061) in 49 ml of neurobasal A medium) at 37 °C in 5% CO_2_. Half of the culture medium (500 µl of 1 ml total volume) was replaced with fresh culture medium every 3 days. At 3 days in vitro (DIV3), 5-fluoro-2’-deoxyuridine (FUDR; Sigma-Aldrich, F0503-100MG), prepared as a 20 mM stock solution in water, was added to the culture at a final concentration of 20 µM to remove non-neuronal cells. On DIV6, the FUDR-containing medium was completely removed and replaced with 500 µl of fresh culture medium. During the medium changes, exposure of the cells to ambient air was minimized. Primary hippocampal neurons were transduced with AAV at DIV7, and transduction efficiency was evaluated at DIV14. These experiments were conducted in accordance with the guidelines of the IACUC at KAIST under approved protocol KA2024-147-v1.

### Neonatal ICV injection and adult RO injection of AAV vectors

All in vivo experiments were conducted in accordance with the guidelines of the IACUC at KAIST under approved protocol KA2025-050-v1. Neonatal ICV injections were performed based on a previously published method^49^ with minor modifications. Postnatal day 2 pups were used, and coordinates for injection ((mediolateral, anteroposterior, dorsoventral) = (0.8, 1.5, 1.5) mm relative to lambda) were controlled using a digital stereotaxic instrument (RWD, 68803) with a custom-made neonatal head holder. For unilateral ICV injections, 5 µl of AAV samples containing 0.1% Fast Green solution was loaded into a 10 µl gastight syringe (HAMILTON, 7653-01) fitted with a 32-gauge small hub removable needle (HAMILTON, 7803-04). Before and during the injection, pups were anesthetized by hypothermia on ice, and the total anesthesia duration did not exceed 5 min. The injection volume was limited to 2 µl for unilateral injection. Immediately after injection, neonatal pups were placed on a 37 °C heating pad for at least 10 min to restore body temperature and were subsequently returned to the dam. Maternal behavior was closely monitored to ensure normal nursing. If nursing behavior was not observed, pups were transferred to a foster dam (ICR; KOATECH).

RO injections were performed in 6-week-old male C57BL/6N mice based on a previously published protocol^50^. AAV samples containing 0.1% Fast Green solution were loaded into a 30-gauge insulin syringe (BD Ultra-Fine, 328868). Anesthesia was induced with 3% isoflurane. Under anesthesia, the left eye was protruded, and the needle was inserted into the retro-orbital sinus with the bevel facing away from the eye. The syringe was slowly advanced, and 100 µl of the AAV samples was injected.

### Tissue collection

All tissues were collected 4 weeks after AAV administration. For fluorescent protein expression analysis, mice were anesthetized with 3% isoflurane and euthanized by transcardial perfusion. Animals were sequentially perfused with 20 ml of PBS followed by 20 ml of 4% PFA. Brains were harvested and post-fixed overnight in 4% PFA at 4 °C, followed by washing in PBS. Fixed tissues were sectioned at 60 µm thickness using a vibrating microtome (Leica Biosystems, VT1200S) and mounted onto glass slides with VECTASHIELD Antifade Mounting Medium with DAPI (Vector LABORATORIES, H-1200). For prime editor expression analysis, mice were euthanized by CO_2_ asphyxiation without perfusion. The cerebral cortex was rapidly dissected, and genomic DNA was extracted using the PureLink Genomic DNA Mini Kit (Thermofisher, K182001) according to the manufacturer’s instructions.

### Imaging analysis for fluorescent protein expression

Fluorescent protein expression in cultured neurons and mouse brain sections was imaged using a Nikon Eclipse Ti-2 confocal microscope equipped with a ×10 air objective lens (numerical aperture 0.45). Quantitative image analysis was performed using ImageJ software. Nuclei were segmented using Otsu’s automatic thresholding method. To separate touching or overlapping nuclei, the Watershed function was applied. Nuclei were quantified using the Analyze Particles function with a size threshold set from 100 pixels to infinity. mNeonGreen- and tdTomato-positive cells were manually counted. Quantification was performed in a blinded manner. To cross-validate the quantification of fluorescence-positive cells in vitro and in in vivo brain slices, additional analyses were performed using Cellpose^51^ (v. 4.0.8). In addition, DAPI-stained nuclei in brain sections were quantified using Cellpose using the pre-trained nuclei model, with automated estimation of the diameter parameter.

### Prime editor efficiency analysis

To evaluate prime editing efficiency in primary hippocampal neurons, genomic DNA was extracted using the EZ Genomic DNA Prep kit (from cultured cells; Enzynomics, EP401-50N) according to the manufacturer’s instructions. For in vivo analysis, neocortical and hippocampal samples were collected, homogenized, and processed using a PureLink Genomic DNA Mini Kit (Invitrogen, K182001) following the manufacturer’s instructions. The genomic region containing the edited *Dnmt1* locus was amplified using Phusion High-Fidelity DNA Polymerase (NEB, M0530L) for 15 PCR cycles to minimize amplification bias. PCR products were purified using an AccuPrep PCR/Gel Purification Kit (Bioneer, K-3037) or DNA Clean & Concentrator-25 kit (Zymo Research, D4033). Quantitative PCR was performed using two primer sets: one targeting the region containing the edited site and another targeting a region unrelated to the editing site as an internal reference. Primer sequences are provided in Supplementary Table 1. Editing efficiency was calculated using the △Ct method by normalizing the edited locus signal to the internal reference locus signal.

### Statistical analysis

Data are presented as mean and s.e.m. Sample size and the statistical tests used for each experiment are described in the figure legends. No statistical methods were used to pre-determine sample size. Statistical analysis was performed using GraphPad Prism software.

## Data availability

The data that support the findings of this study are available in the paper and its Supplementary Information. pENN.AAV.EF1a.eGFP.WPRE.rBG, pAAV2/9n, pAdDeltaF6, pAAV-CAG-tdTomato, pAAV-CAG-2xNLS-tdTomato-WPRE-hGHpolyA, v3em-Cterm-PE6d-dualU6-Dnmt1-PE3-loxp, v3em-Nterm-PE2max and pUCmini-iCAP-AAV.CAP-B10 plasmids are available from Addgene pending scientific review and a completed material transfer agreement.

## Acknowledgements

We thank Catherine Oikonomou for help with manuscript editing. We thank Seongmin Jang, Tae Kyoung Lee, and Gerard M. Coughlin for helpful discussions. We thank James M. Wilson, Edward Boyden, Guoping Feng, and David Liu for the generous gifts of the plasmids. Figures were created using BioRender.com. The work was supported by National Research Foundation of Korea (NRF) grants funded by the Korean government (Ministry of Science and ICT, MSIT) (RS-2023-00210139, RS-2024-00436904, RS-2024-00401376) to J.P. V.G. is an Investigator of the Howard Hughes Medical Institute.

## Author information

Y.K., V.G., and J.P. designed the study. Y.K., X.D., and J.P. developed the AAV linking strategy. Y.K. performed molecular cloning, viral production, AAV linking, western blot, in vitro experiments, mouse injections, and prime editing analysis. Y.K., G.L., and C.Y. performed perfusion, imaging, and data analysis. Y.K. and J.P. prepared figures and wrote the manuscript. X.D. and V.G. reviewed the manuscript. V.G. and J.P. supervised all aspects of the study.

## Competing interests

The authors declare no competing interests.

## Notes

### Competing Interest Statement

The authors have declared no competing interest.

## References

1. Ling, Q., Herstine, J.A., Bradbury, A. & Gray, S.J. AAV-based in vivo gene therapy for neurological disorders. Nat.Rev.Drug.Discov. 22, 789–806 (2023).

2. Haery, L. & Fan, M. Adeno-Associated Virus Technologies and Methods for Targeted Neuronal Manipulation. Front.Neuroanat. 13, 93 (2019).

3. Costa Verdera, H., Kuranda, K. & Mingozzi, F. AAV Vector Immunogenicity in Humans: A Long Journey to Successful Gene Transfer. Mol.Ther. 28, 723–746 (2020).

4. Wu, X., Jiang, Y., Pu, K. & Hong, G. Tether-free photothermal deep-brain stimulation in freely behaving mice via wide-field illumination in the near-infrared-II window. *Nat*. Biomed. Eng 6, 754–770 (2022).

5. Wu, Z., Asokan, A. & Samulski, R.J. Adeno-associated virus serotypes: vector toolkit for human gene therapy. Mol.Ther. 14, 316–327 (2006).

6. Nelson, C.E. & Gersbach, C.A. Long-term evaluation of AAV-CRISPR genome editing for Duchenne muscular dystrophy. Nat.Med. 25, 427–432 (2019).

7. Mich, J.K. & Kalume, F.K. Interneuron-specific dual-AAV SCN1A gene replacement corrects epileptic phenotypes in mouse models of Dravet syndrome. Sci.Transl.Med. 17, eadn5603 (2025).

8. Lindley, S.R. & Anderson, D.M. Ribozyme-activated mRNA trans-ligation enables large gene delivery to treat muscular dystrophies. Science 386, 762–767 (2024).

9. Yang, Y. & Wilson, J.M. A dual AAV system enables the Cas9-mediated correction of a metabolic liver disease in newborn mice. Nat.Biotechnol. 34, 334–338 (2016).

10. Davis, J.R. & Liu, D. Efficient prime editing in mouse brain, liver and heart with dual AAVs. Nat.Biotechnol. 42, 253–264 (2023).

11. Doman, J.L., Pandey, S. & Liu, D. Phage-assisted evolution and protein engineering yield compact, efficient prime editors. Cell 186, 3983–4002.e3926 (2023).

12. Tabebordbar, M. et al. In vivo gene editing in dystrophic mouse muscle and muscle stem cells. Science 351, 407–411 (2016).

13. Nelson, C.E. et al. In vivo genome editing improves muscle function in a mouse model of Duchenne muscular dystrophy. Science 351, 403–407 (2016).

14. Riedmayr, L.M. & Becirovic, E. mRNA trans-splicing dual AAV vectors for (epi)genome editing and gene therapy. Nat.Commun. 14, 6578 (2023).

15. Tornabene, P. & Auricchio, A. Intein-mediated protein trans-splicing expands adeno-associated virus transfer capacity in the retina. Sci.Transl.Med. 11, 492 (2019).

16. Duan, D., Yue, Y. & Engelhardt, J.F. Expanding AAV packaging capacity with trans-splicing or overlapping vectors: a quantitative comparison. Mol.Ther. 4, 383–391 (2001).

17. Yan, Z., Zhang, Y., Duan, D. & Engelhardt, J.F. Trans-splicing vectors expand the utility of adeno-associated virus for gene therapy. Proc.Natl.Acad.Sci.USA 97, 6716–6721 (2000).

18. Johnston, S., Parylak, S.L., Kim, S. & Shtrahman, M. AAV ablates neurogenesis in the adult murine hippocampus. Elife 10 (2021).

19. Stone, D., Aubert, M. & Jerome, K.R. Adeno-associated virus vectors and neurotoxicity-lessons from preclinical and human studies. Gene.Ther. 32, 60–73 (2025).

20. Colella, P., Ronzitti, G. & Mingozzi, F. Emerging Issues in AAV-Mediated In Vivo Gene Therapy. Mol.Ther.Methods.Clin.Dev. 8, 87–104 (2018).

21. Nagao, K. & Anikeeva, P. Adeno-associated viruses escort nanomaterials to specific cells and tissues. Preprint at bioRxiv 10.1101/2025.04.04.647267 (2025).

22. Byrne, L., Ozturk, B.E. & Day, T., Modular system for gene and protein delivery based on aav. US patent 20200407751A1 (2025).

23. Liu, W., Halverson, J., Tian, Y., Tkachenko, A.V. & Gang, O. Self-organized architectures from assorted DNA-framed nanoparticles. Nat.Chem. 8, 867–873 (2016).

24. Maye, M.M., Nykypanchuk, D., Cuisinier, M., Van Der Lelie, D. & Gang, O. Stepwise surface encoding for high-throughput assembly of nanoclusters. Nat.Mater. 8, 388–391 (2009).

25. Cui, H. & Wei, B. Stepwise Assembly of DNA Nanostructures in a Surface-Based Method. ACS Nano 18, 31773–31779 (2024).

26. Chan, M.S. & Lo, P.K. Stepwise Ligand-induced Self-assembly for Facile Fabrication of Nanodiamond-Gold Nanoparticle Dimers via Noncovalent Biotin-Streptavidin Interactions. Nano Lett. 19, 2020–2026 (2019).

27. Zhang, D.Y. & Winfree, E. Control of DNA strand displacement kinetics using toehold exchange. J.Am.Chem.Soc. 131, 17303–17314 (2009).

28. Blizzard, R. J., Gibson, T. E., Mehl, R. A. Site-specific protein labeling with tetrazine amino acids. Methods Mol. Biol. 1728, 201–217 (2018).

29. Kim, E. & Koo, H. Biomedical applications of copper-free click chemistry:: In vitro, in vivo, and ex vivo. Chem.Sci. 10, 7835–7851 (2019).

30. Oliveira, B.L., Guo, Z. & Bernardes, G.J.L. Inverse electron demand Diels-Alder reactions in chemical biology. Chem.Soc.Rev. 46, 4895–4950 (2017).

31. Seo, J.W. & Ferrara, K.W. Positron emission tomography imaging of novel AAV capsids maps rapid brain accumulation. Nat.Commun. 11 (2020).

32. Knall, A.C. & Slugovc, C. Inverse electron demand Diels-Alder (iEDDA)-initiated conjugation: a (high) potential click chemistry scheme. Chem.Soc.Rev. 42, 5131–5142 (2013).

33. Lim, C.Y. & Porter, M.D. Succinimidyl ester surface chemistry: implications of the competition between aminolysis and hydrolysis on covalent protein immobilization. Langmuir 30, 12868–12878 (2014).

34. Zadeh, J.N., Steenberg, C.D., Bois, J.S., Wolfe, B.R. & Pierce, N.A. NUPACK: Analysis and design of nucleic acid systems. J.Comput.Chem. 32, 170–173 (2011).

35. Shchepinov, M.S., Case-Green, S.C. & Southern, E.M. Steric factors influencing hybridisation of nucleic acids to oligonucleotide arrays. Nucleic.Acids.Res. 25, 1155–1161 (1997).

36. Long, D. & Tian, Z. Understanding the relationship between sequences and kinetics of DNA strand displacements. Nucleic.Acids.Res. 52, 9407–9416 (2024).

37. Hughes, M.P. & Rahim, A.A. AAV9 intracerebroventricular gene therapy improves lifespan, locomotor function and pathology in a mouse model of Niemann-Pick type C1 disease. Hum.Mol.Genet. 27, 3079–3098 (2018).

38. Gonzalez, T.J. & Asokan, A. Cross-species evolution of a highly potent AAV variant for therapeutic gene transfer and genome editing. Nat.Commun. 13, 5947 (2022).

39. Park, J., O’Shea, H. & Lee, S. AAV-mediated neuronal expression of FOXG1 restores oligodendrocyte maturation, myelination, and hippocampal structure in mouse models of FOXG1 syndrome. Preprint at bioRxiv 10.64898/2025.12.04.692422 (2025).

40. Aoki, R., Konno, A., Hosoi, N., Kawabata, H. & Hirai, H. AAV vectors for specific and efficient gene expression in microglia. Cell.Rep.Methods 5, 101116 (2025).

41. Chuapoco, M.R., Flytzanis, N.C., Goeden, N. & Gradinaru, V. Adeno-associated viral vectors for functional intravenous gene transfer throughout the non-human primate brain. Nat. Nanotechnol. 18, 1241–1251 (2023).

42. Jang, S., Shen, H.K., Ding, X., Miles, T.F. & Gradinaru, V. Structural basis of receptor usage by the engineered capsid AAV-PHP.eB. Mol.Ther.Methods.Clin.Dev. 26, 343–354 (2022).

43. Brittain, T.J., Jang, S. & Gradinaru, V. Structural basis of liver de-targeting and neuronal tropism of CNS-targeted AAV capsids. Preprint at bioRxiv 10.1101/2025.06.02.655683 (2025).

44. Blizzard, R.J. & Mehl, R.A. Ideal Bioorthogonal Reactions Using A Site-Specifically Encoded Tetrazine Amino Acid. J.Am.Chem.Soc. 137, 10044–10047 (2015).

45. Wang, Y. & Liu, T. Noncanonical amino acids as doubly bio-orthogonal handles for one-pot preparation of protein multiconjugates. Nat.Commun. 14, 974 (2023).

46. Liu, W., Brock, A., Chen, S., Chen, S. & Schultz, P.G. Genetic incorporation of unnatural amino acids into proteins in mammalian cells. Nat.Methods 4, 239–244 (2007).

47. Challis, R.C., Kumar, S.R. & Gradinaru, V. Systemic AAV vectors for widespread and targeted gene delivery in rodents. Nat.Protoc. 14, 379–414 (2019).

48. Cramer, T.M.L. & Tyagarajan, S.K. Protocol for the culturing of primary hippocampal mouse neurons for functional in vitro studies. STAR.Protoc. 5, 102991 (2024).

49. Yang, K., Li, T. & Li, F. Protocol for the neonatal intracerebroventricular delivery of adeno-associated viral vectors for brain restoration of MECP2 for Rett syndrome. STAR.Protoc. 5, 103344 (2024).

50. Yardeni, T., Eckhaus, M., Morris, H. D., Huizing, M., & Hoogstraten-Miller, S. Retro-orbital injections in mice. Lab.Anim. 40, 155–160 (2011).

51. Stringer, C., Wang, T., Michaelos, M. & Pachitariu, M. Cellpose: a generalist algorithm for cellular segmentation. Nat.Methods. 18, 100–106 (2021).

