## Supplementary Information for "Controlled linking of AAV capsids enables coordinated multi-vector delivery"

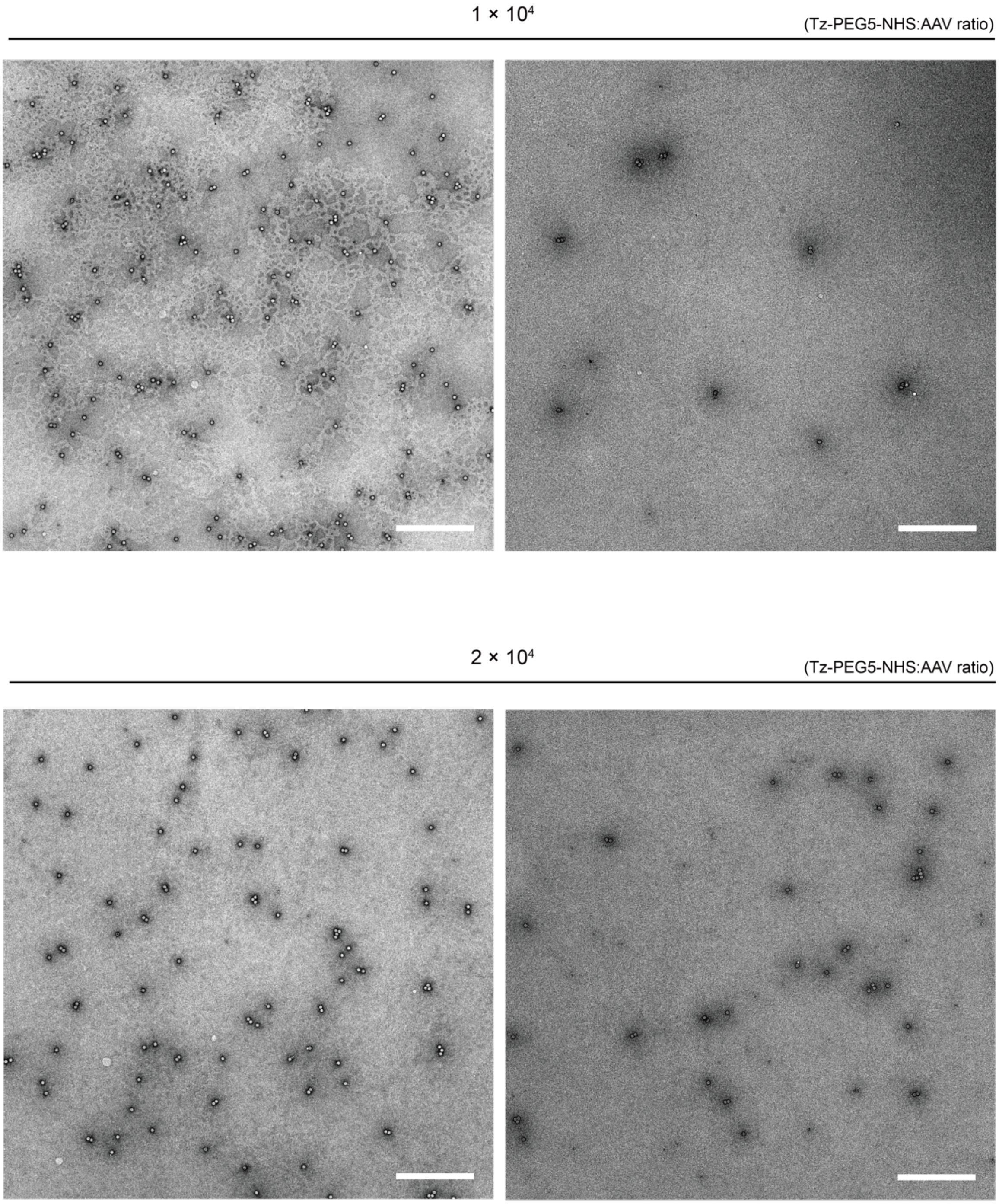


**Supplementary Figure 1.** Representative low-magnification TEM images of linked AAVs prior to purification at Tz-PEG5-NHS:AAV ratios of $1.0 \times 10^4$ and $2.0 \times 10^4$. Quantification of AAV monomer, dimer, trimer, and higher-order aggregate populations is presented in Fig. 2e (scale bars, 500 nm).

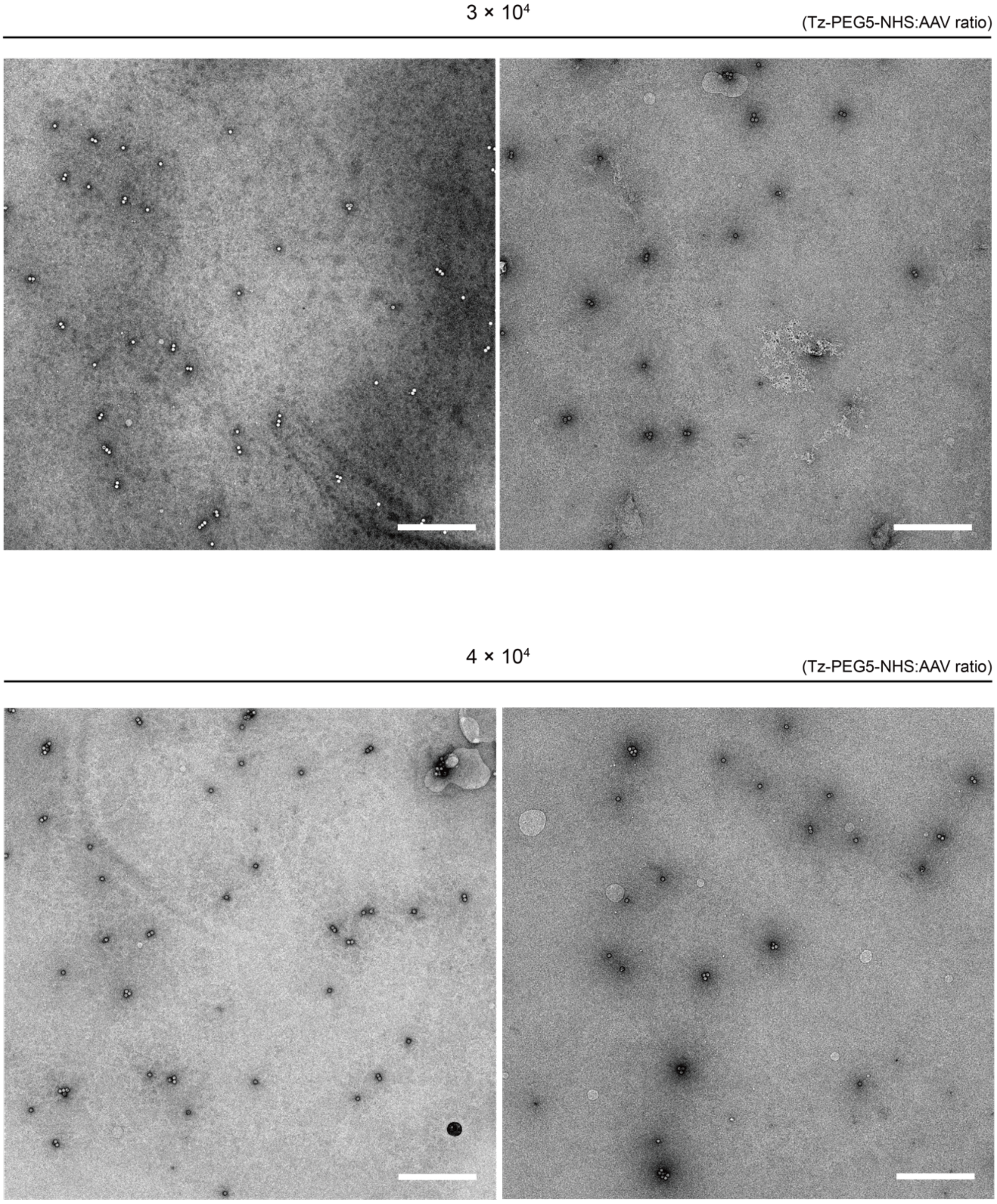


**Supplementary Figure 2.** Representative low-magnification TEM images of linked AAVs prior to purification at Tz-PEG5-NHS:AAV ratios of $3.0 \times 10^4$ and $4.0 \times 10^4$. Quantification of AAV monomer, dimer, trimer, and higher-order aggregate populations is presented in Fig. 2e (scale bars, 500 nm).

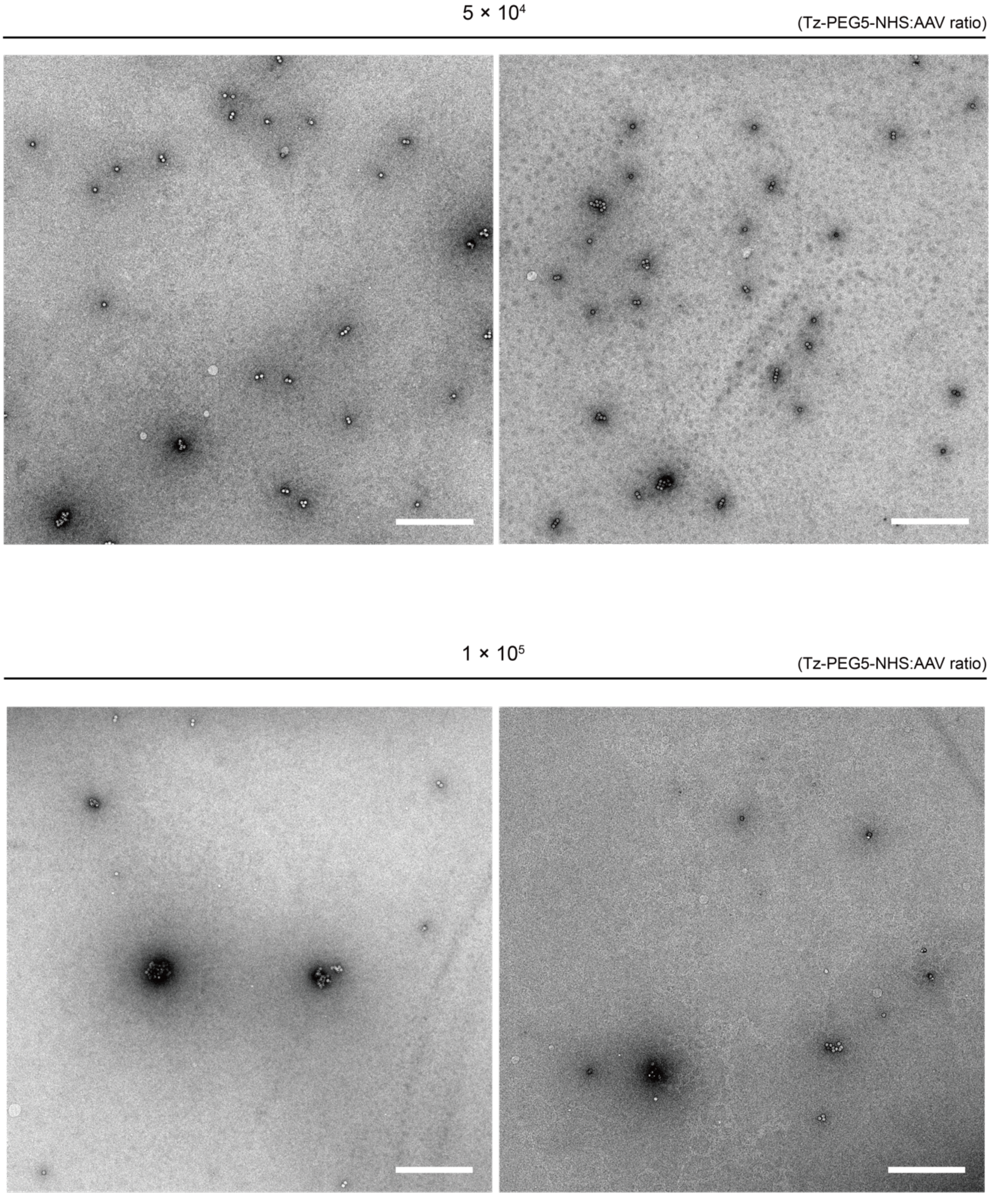


**Supplementary Figure 3.** Representative low-magnification TEM images of linked AAVs prior to purification at Tz-PEG5-NHS:AAV ratios of $5.0 \times 10^4$ and $1.0 \times 10^5$. Quantification of AAV monomer, dimer, trimer, and higher-order aggregate populations is presented in Fig. 2e. As the Tz–PEG5–NHS:AAV ratio increased, the formation of higher-order aggregates became more prominent, and the number of capsids per aggregate increased (scale bars, 500 nm).

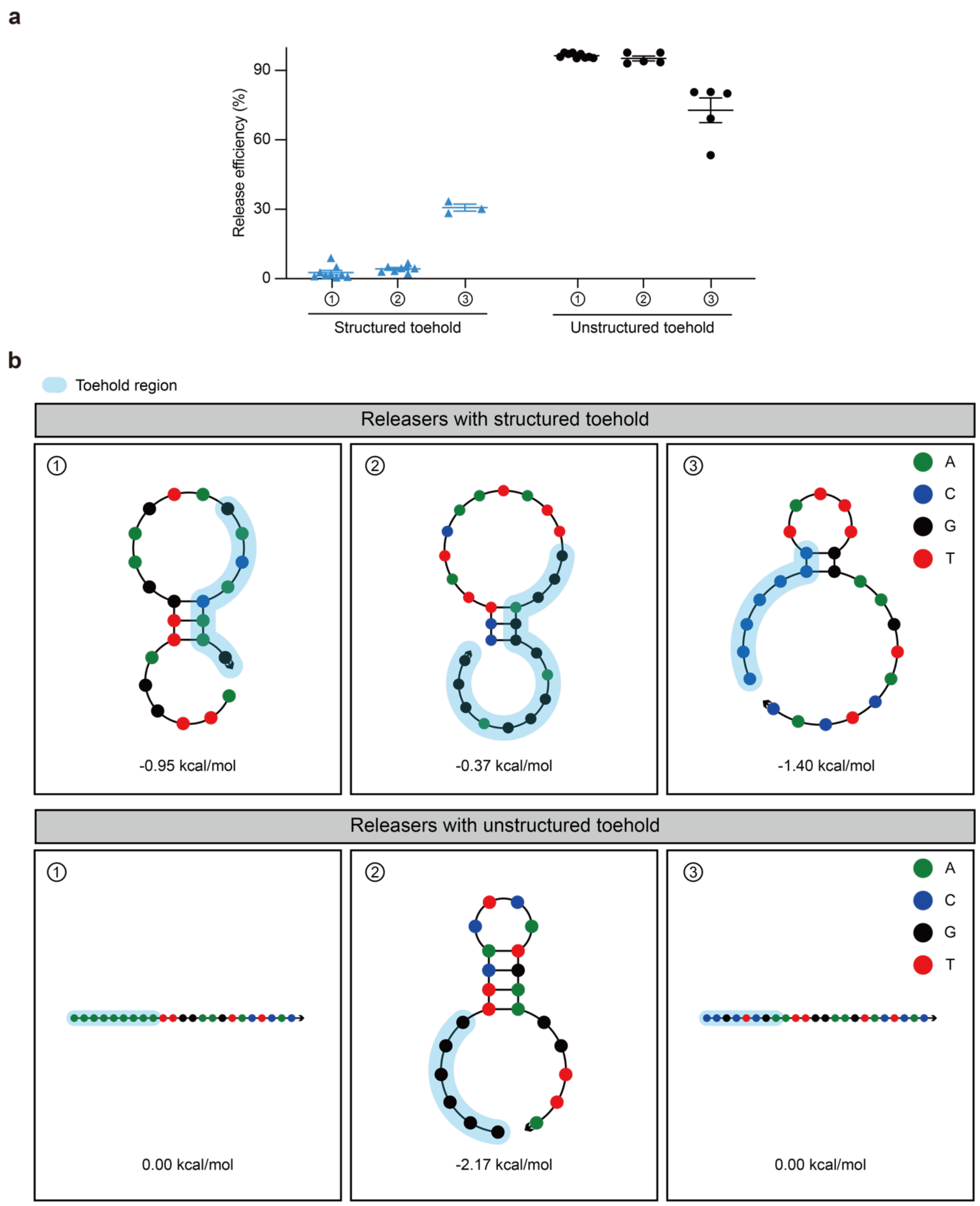


**Supplementary Figure 4.** Effects of toehold structure on the release of surface-bound AAVs. **(a)** Fraction of released AAVs after 3 h incubation with releaser strands containing either structured or unstructured toehold regions. **(b)** Sequences and predicted secondary structures of the releaser strands evaluated in (a). Secondary structures were predicted using NUPACK at

25 °C under ionic conditions of 300 mM $Na^+$ and 12.5 mM $Mg^{2+}$, mimicking the reaction conditions for the release of surface-bound AAVs. The predicted free energy values (ΔG, kcal $mol^{-1}$) are shown below each structure. For (a), data are presented as mean ± s.e.m. from at least three independent experiments.

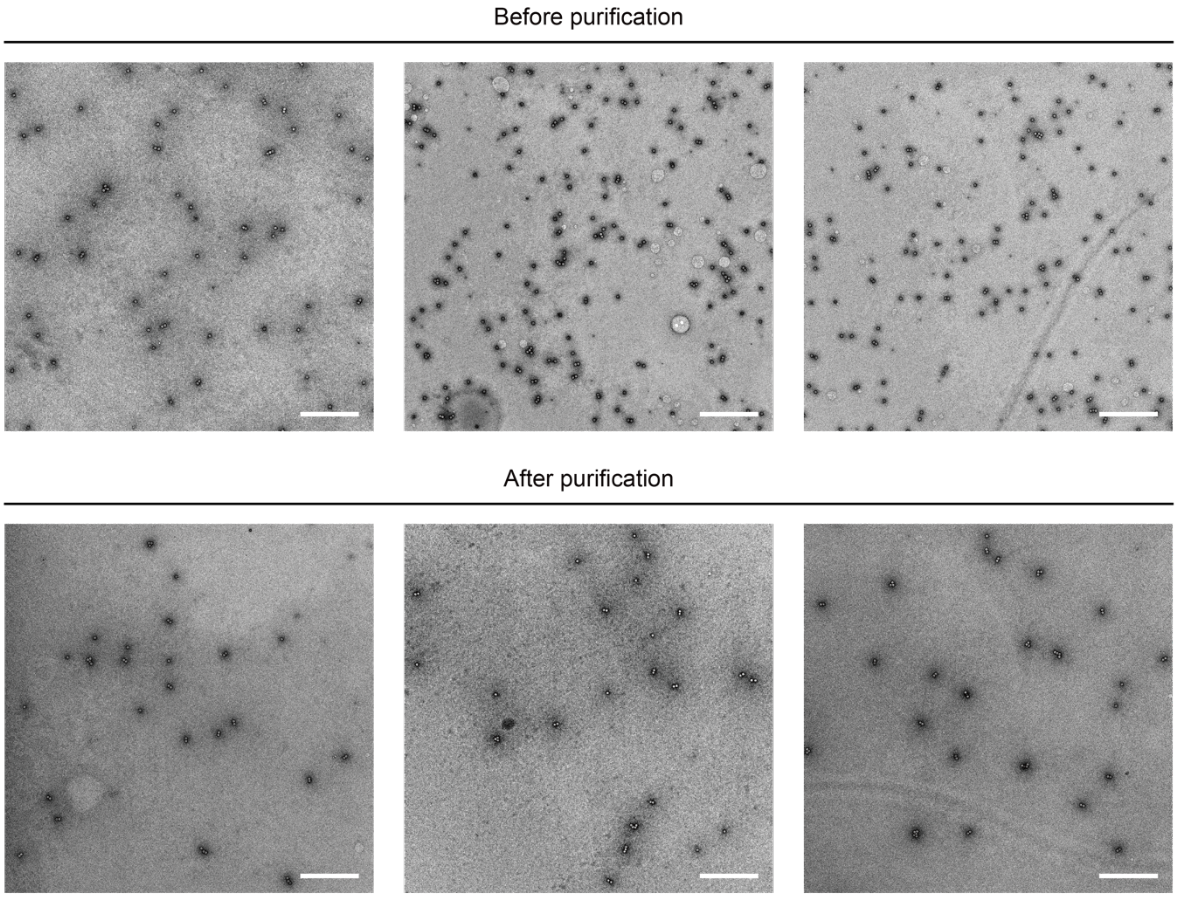


**Supplementary Figure 5.** Representative low-magnification TEM images of linked AAVs before and after the purification step. A marked decrease in the monomer population is clearly observed after purification. The distribution of AAV monomer, dimer, trimer, and higher-order aggregate populations before and after purification is quantified in Fig. 2h (scale bars, 500 nm).

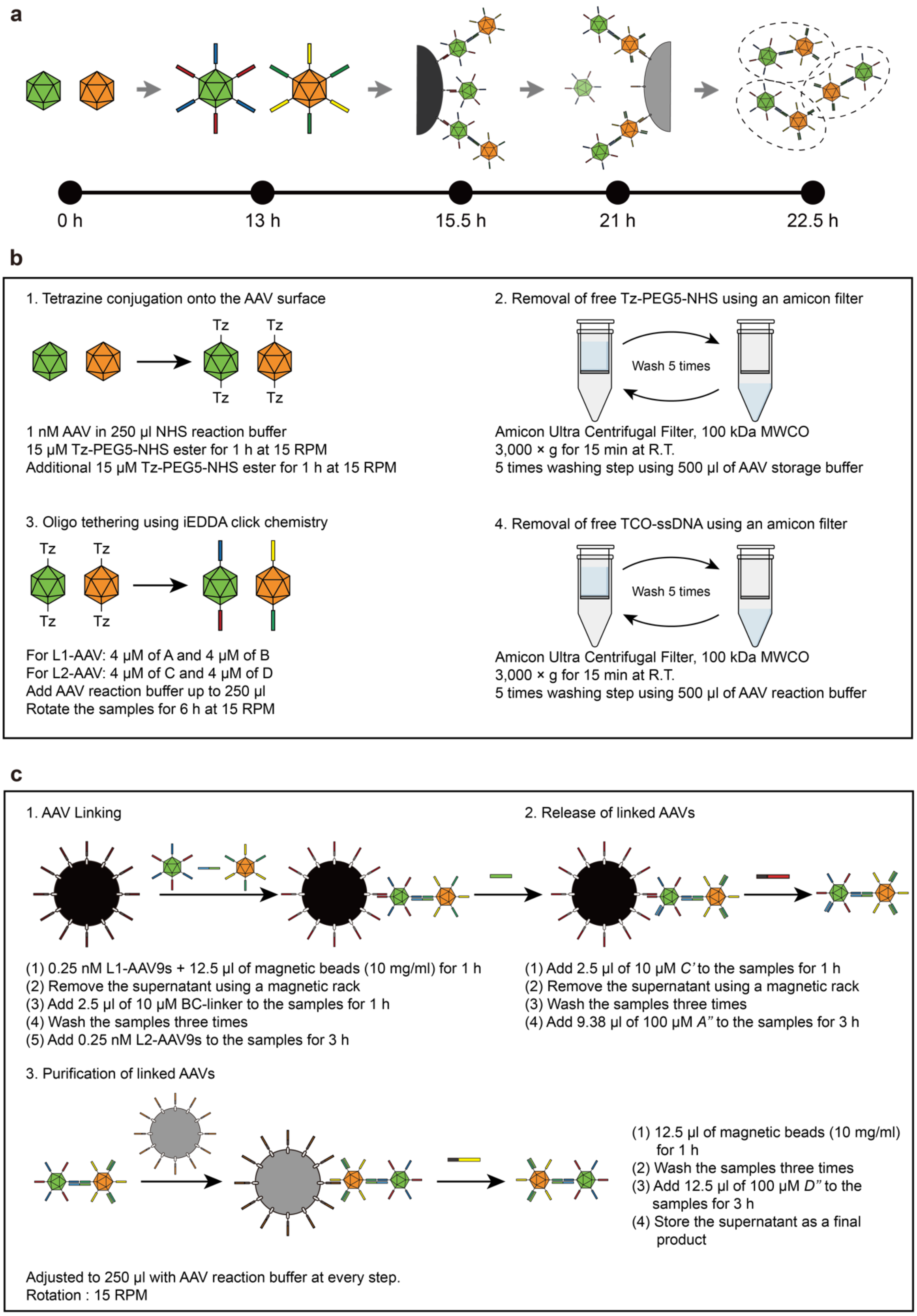


**Supplementary Figure 6.** Processing time for the generation of linked AAVs. **(a)** A schematic overview of the complete workflow, including oligonucleotide conjugation, L1-AAV anchoring, inter-AAV linking, toehold-mediated release, and purification, with associated processing times for each step. **(b)** Detailed protocol and processing time for oligonucleotide conjugation onto the AAV capsid surface via iEDDA click chemistry. **(c)** Detailed protocol and processing time for L1-AAV anchoring, inter-AAV linking, toehold-mediated release of surface-bound AAVs, and purification of linked AAVs.

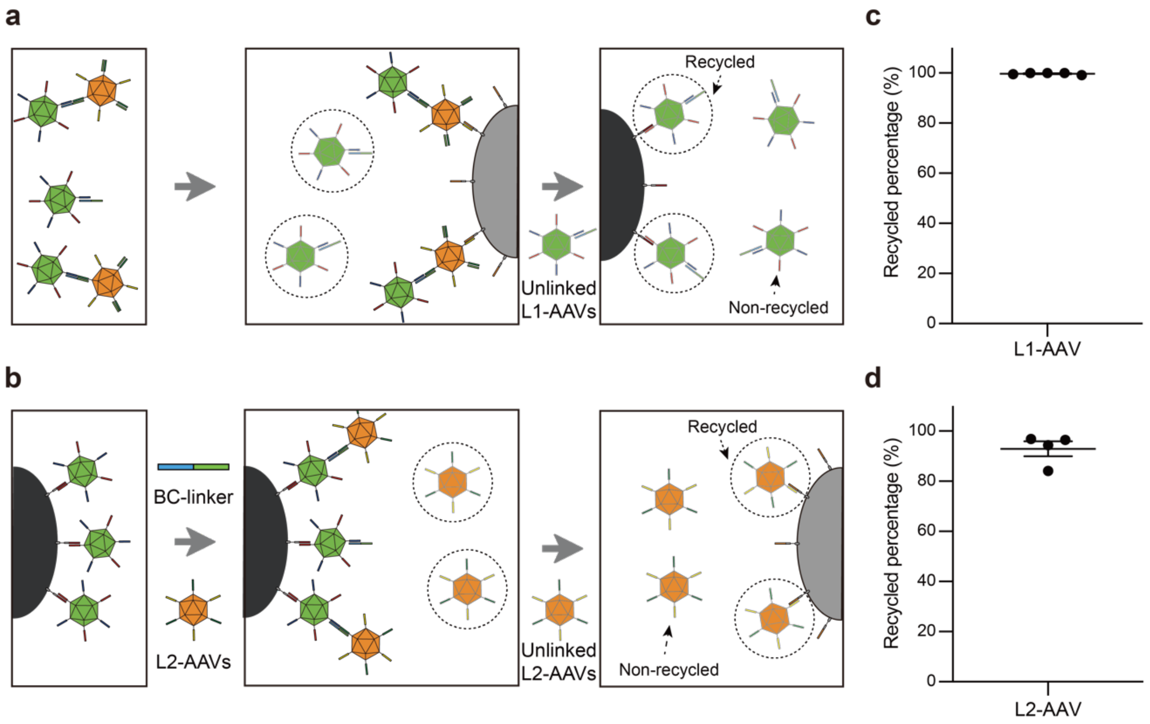


**Supplementary Figure 7.** Recycling of unlinked AAVs. **(a, b)** Schematic diagrams illustrating the experimental workflows used to evaluate the recovery and reuse of unlinked L1-AAVs (a) and L2-AAVs (b), respectively. **(c, d)** qPCR-based AAV titration quantifying the percentage of recovered versus unrecovered L1-AAVs (c) and L2-AAVs (d), respectively.

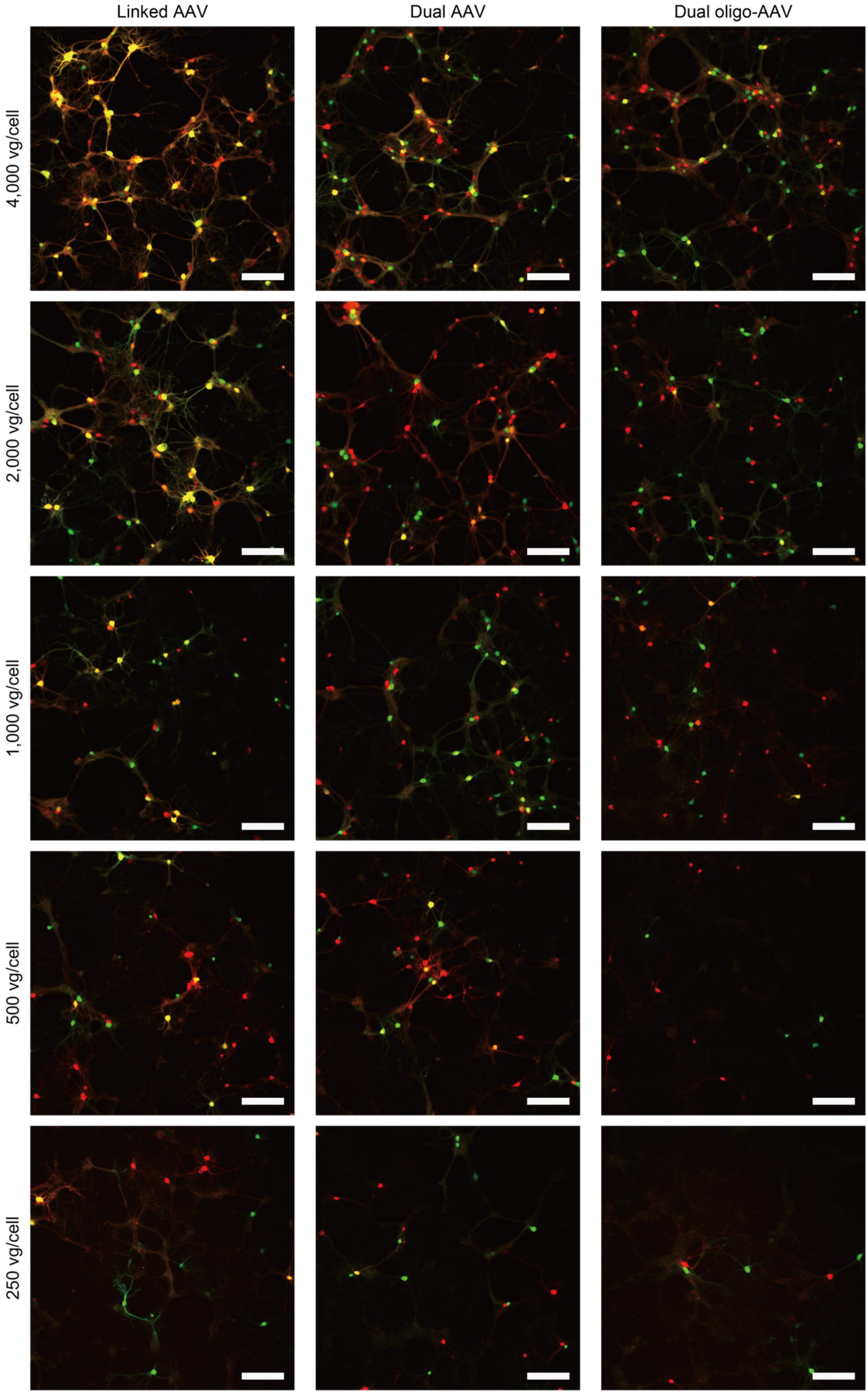
Linked AAV
Dual AAV
Dual oligo-AAV
4,000 vg/cell
2,000 vg/cell
1,000 vg/cell
500 vg/cell
250 vg/cell

**Supplementary Figure 8.** Representative fluorescence images of primary hippocampal neurons transduced with linked AAV, dual AAV, or dual oligo-AAV at doses ranging from 4,000 (top) to 250 (bottom) vg/cell. Red fluorescence represents transduction by L2-AAV9 (AAV9-CAG::tdTomato), whereas green fluorescence indicates transduction by L1-AAV9 (AAV9-CAG::mNeonGreen). Fluorescence images were acquired under identical imaging settings (scale bars, 200 μm).

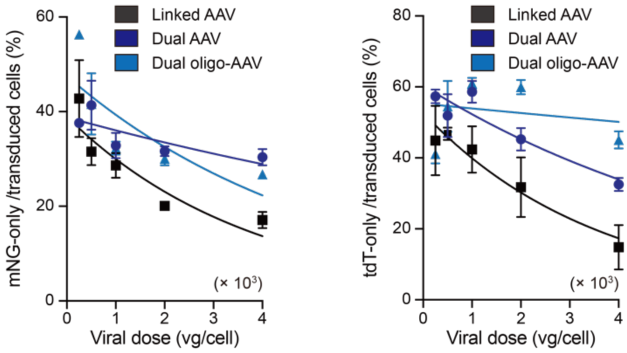


**Supplementary Figure 9.** Quantification of neurons expressing only mNeonGreen or only tdTomato across viral doses. Percentages were calculated as the number of neurons expressing mNeonGreen only (left) or tdTomato only (right) divided by the total number of transduced neurons within each ROI and multiplied by 100. Data represent mean ± s.e.m. from at least three independent biological replicates.

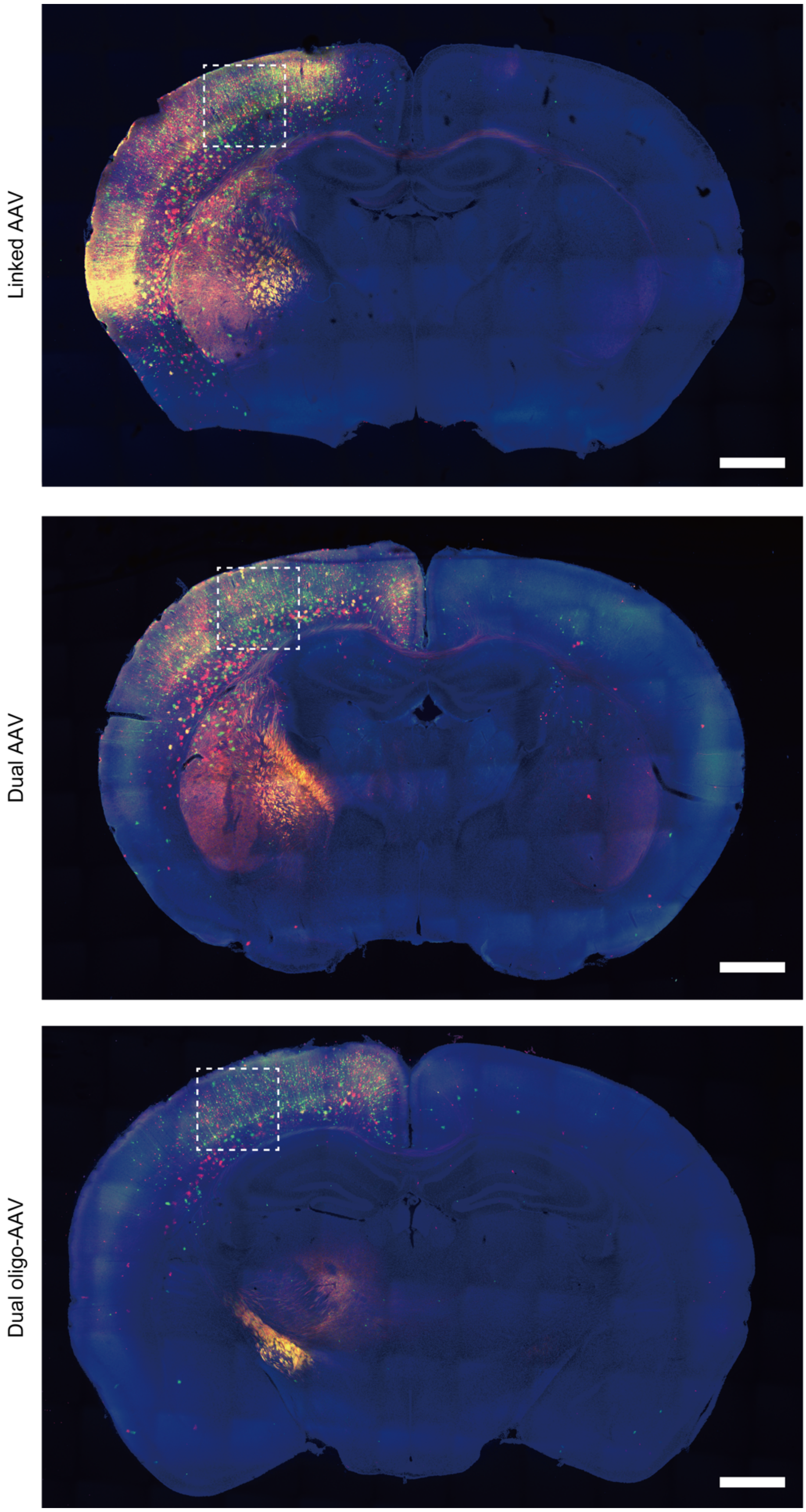


**Supplementary Figure 10.** Representative whole-brain fluorescence images following ICV delivery of linked AAVs, dual AAVs, or dual oligo-AAVs at a dose of $2.5 \times 10^8$ vg per pup. Red fluorescence indicates transduction by L2-AAV9 (AAV9-CAG::tdTomato), and green fluorescence represents transduction by L1-AAV9 (AAV9-CAG::mNeonGreen). Strong fluorescent signals were observed across neocortex. Representative ROIs used for the analysis in Fig. 3h-i are outlined with dashed rectangles. Images were acquired under identical imaging settings (scale bars, 1,000 μm).

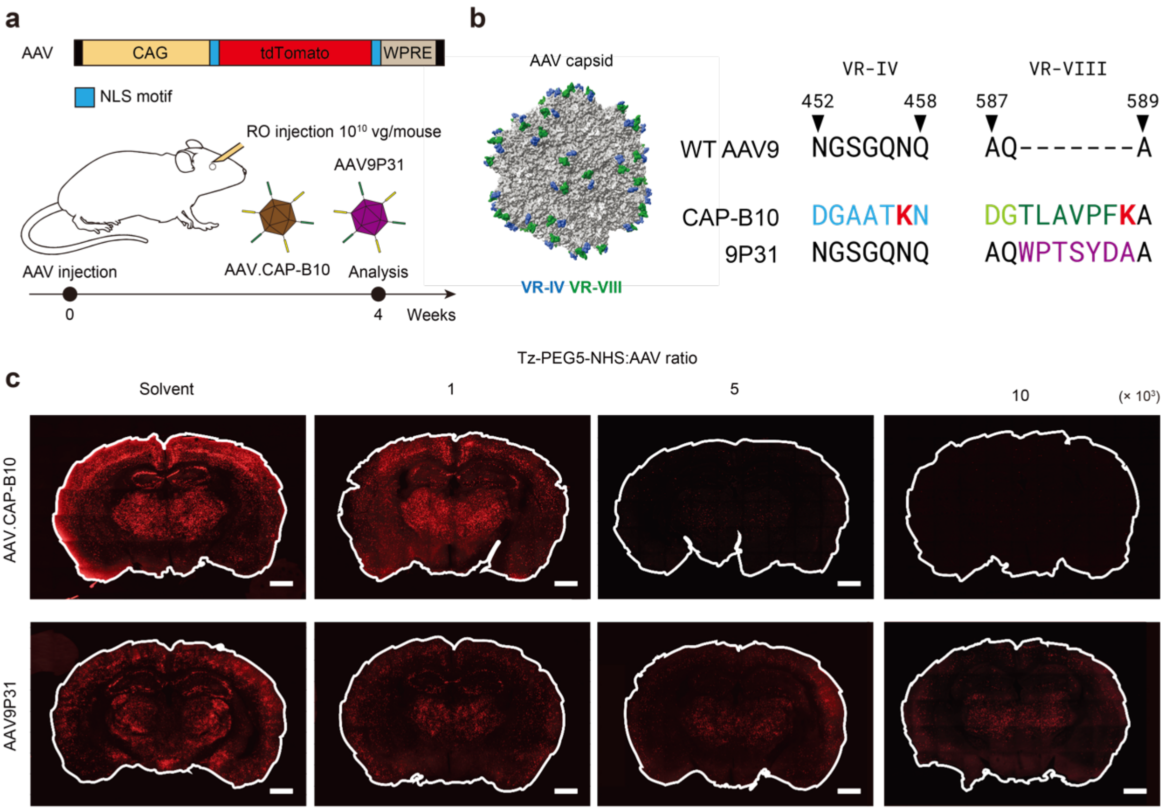


**Supplementary Figure 11.** Effects of oligonucleotide conjugation on the tropism of engineered AAV capsids. **(a)** In vivo experimental scheme for systemic RO injection of oligonucleotide-functionalized engineered AAV variants (AAV.CAP-B10 and AAV9P31), followed by whole-brain analysis four weeks post-injection. **(b)** Sequence alignment of VR-IV and VR-VIII regions from CAP-B10 and 9P31. Lysine residues are highlighted in bold and red. **(c)** Representative whole-brain fluorescence images showing differential preservation of brain tropism following oligonucleotide functionalization (scale bars, 1,000 μm). Increasing Tz–PEG5–NHS:AAV ratios progressively reduced brain transduction of AAV.CAP-B10, whereas AAV9P31 largely retained its engineered tropism, indicating capsid-dependent tolerance to surface modification.

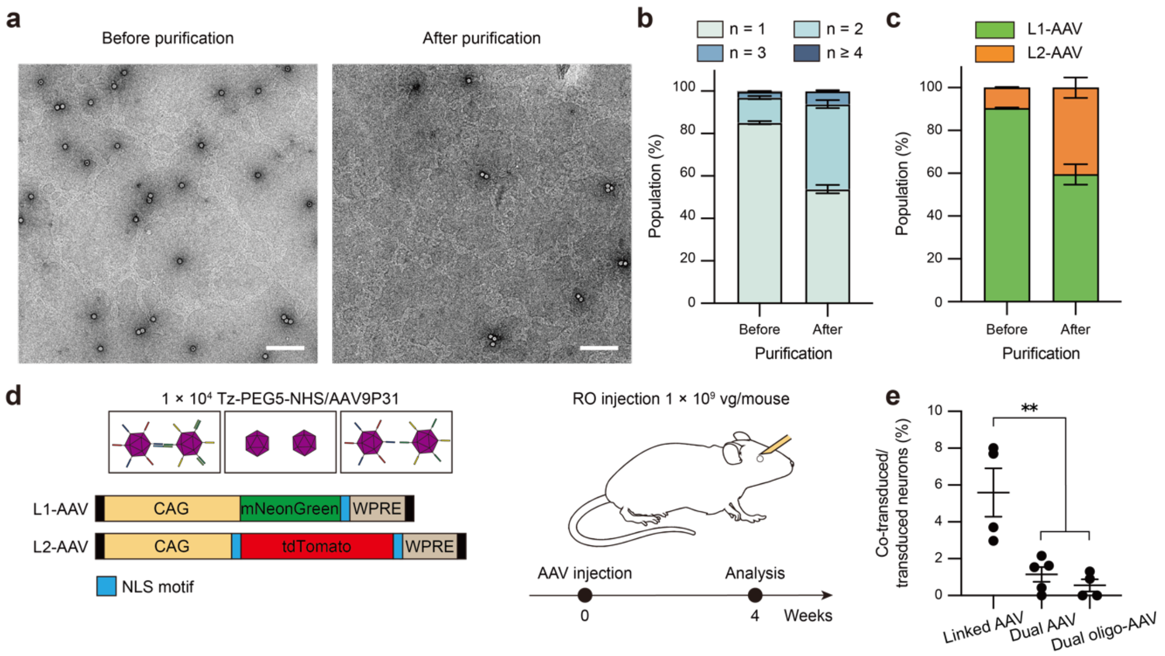


**Supplementary Figure 12.** Systemic co-delivery via linked AAV9P31. **(a)** Representative TEM images of linked AAV9P31 before and after purification (scale bars, 200 nm). AAV9P31 was modified at a Tz-PEG5-NHS:AAV ratio of $1.0 \times 10^4$ for the linking. **(b)** Distribution of monomers, dimers, and higher-order linked AAV9P31 before and after purification based on TEM results. **(c)** Genome composition of linked AAV9P31 before and after purification. **(d)** Schematic of linked AAV9P31 and control groups (dual AAV9P31 and dual oligo-AAV9P31) encoding mNeonGreen and tdTomato, along with the experimental timeline for in vivo RO injection at $1 \times 10^9$ vg per mouse. **(e)** Quantification of expression uniformity in striatum ROIs four weeks post-injection. Graphs in (b), (c), and (e) represent mean ± s.e.m. from at least three independent biological replicates. For (e), statistical analysis was performed using one-way ANOVA. Statistical significance is indicated as follows: $^{**}p < 0.01$.
